# Brain Tumor IDH, 1p/19q, and MGMT Molecular Classification Using MRI-based Deep Learning: Effect of Motion and Motion Correction

**DOI:** 10.1101/2020.06.01.126375

**Authors:** Sahil S. Nalawade, Fang F. Yu, Chandan Ganesh Bangalore Yogananda, Gowtham K. Murugesan, Bhavya R. Shah, Marco C. Pinho, Benjamin C. Wagner, Bruce Mickey, Toral R. Patel, Baowei Fei, Ananth J. Madhuranthakam, Joseph A. Maldjian

## Abstract

Deep learning has shown promise for predicting glioma molecular profiles using MR images. Before clinical implementation, ensuring robustness to real-world problems, such as patient motion, is crucial. We sought to evaluate the effects of motion artifact on glioma marker classifier performance and develop a deep learning motion correction network to restore classification accuracies. T2w images and molecular information were retrieved from the TCIA and TCGA databases. Three-fold cross-validation was used to train and test the motion correction network on artifact-corrupted images. We then compared the performance of three glioma marker classifiers (IDH mutation, 1p/19q codeletion, and MGMT methylation) using motion-corrupted and motion-corrected images. Glioma marker classifier performance decreased markedly with increasing motion corruption. Applying motion correction effectively restored classification accuracy for even the most motion-corrupted images. For IDH classification, an accuracy of 99% was achieved, representing a new benchmark in non-invasive image-based IDH classification and exceeding the original performance of the network. Robust motion correction can enable high accuracy in deep learning MRI-based molecular marker classification rivaling tissue-based characterization.

**STATEMENT OF SIGNIFICANCE:** Deep learning networks have shown promise for predicting molecular profiles of gliomas using MR images. We demonstrate that patient motion artifact, which is frequently encountered in the clinic, can significantly impair the performance of these algorithms. The application of robust motion correction algorithms can restore the performance of these networks, rivaling tissue-based characterization.

## 1. Introduction

Primary brain neoplasms represent a group of tumors with broad variations in imaging features, response to therapy, and prognosis. It has become evident that the observed clinical heterogeneity is associated with specific molecular and genetic profiles. For example, isocitrate dehydrogenase 1 and 2 (IDH 1/2) mutated gliomas (1) demonstrate significantly increased survival compared to wild-type gliomas with the same histologic grade. Additionally, 1p/19q codeletion(2) and O6-methyl guanine-DNA methyltransferase (MGMT) promoter methylation(3) have been associated with differences in response to specific chemoradiation regimens. These observations led the World Health Organization to revise its classification of gliomas in 2016 (4). In clinical practice, the only way to identify these molecular profiles has been through immunohistochemistry or gene sequencing, requiring surgical tissue retrieval using invasive biopsy or tumor resection. However, obtaining these tissue samples can be challenging, as a report from The Cancer Genome Atlas (TCGA) indicates only 35% of biopsies obtained sufficient tumor for IDH testing (5). Recently, there have been advances in classifying tumor profiles using non-invasive imaging, particularly MR imaging (6,7). The algorithms used for tumor classification can be designed based on linear regression models, classical machine learning (8-10), and, more recently, deep learning networks (11).

Deep learning-based methods have shown particular promise, outperforming other approaches, including classical machine learning. Using tumor maps and molecular labels as ground truth(12,13), the imaging features that help classify the tumor molecular subtype are learned by the algorithm using convolutional layers. Chang et al. recently applied a form of deep-learning known as a convolutional neural network (CNN) to multiparametric MR images and reported accuracies of 94%, 92%, and 83% for IDH mutation status, 1p/19q codeletion, and MGMT methylation, respectively (14). Our group recently achieved 97% accuracy for classifying IDH mutation status in primary brain tumors utilizing T2-weighted (T2w) MR images alone (7). We have extended this approach with T2w images to 1p/19q (15) and MGMT, achieving accuracies of 93% and 95%, respectively, rivaling tissue characterization. Although these preliminary results have been impressive, additional work is needed before these methods can be fully adopted in the clinic.

An important caveat is that the effects of degradation on the input MR images, such as motion artifact, and in turn, on the performance of deep learning-based classifiers, has not been systematically studied. Motion artifacts are an especially pervasive source of MR image quality degradation and can be due to gross patient movements, as well as physiologic cardiac and respiratory motion (16,17). In clinical practice, these artifacts can interfere with diagnostic interpretation, impacting image quality in 10-42% of brain MR examinations (18), and necessitating repeat imaging in up to 20% of cases (19). It is also not guaranteed that a patient will be better able to hold motionless during repeat imaging, and often the diagnostic quality remains impaired. This can incur substantial financial costs to the health care system. Pei et al.(20) applied physical models of motion blurring to non-medical image data and tested the classification performance of two CNNs, including AlexNet and VGGnet, and showed decreased accuracy in classifying the distorted images. It is, therefore, likely that motion corruption will also lead to reduced performance of deep learning algorithms in classifying brain tumor images.

One potential solution is to directly correct or prevent motion artifacts, with multiple methods proposed by the MR research community. These include prospective and retrospective navigator-based approaches, which extract the patient’s positional information from the MR scanner (21,22), as well as the application of motion robust acquisitions, such as the PROPELLER technique (23), which prospectively oversamples the central portions of k-space. However, these methods generally lead to increased scan times and produce residual artifacts even after correction (18). To date, a handful of studies have applied deep learning for the correction of motion artifacts. Duffy et al. (24) utilized a generative adversarial network (GAN) based motion correction algorithm to remove motion artifact in T1w MR brain images obtained from the Autism Brain Imaging Data Exchange database. Fantini et al. (25) proposed a network for automated detection of motion artifact in MR images by training along the axial, coronal, and sagittal axes of the image. Sommer et al. similarly utilized fully convolutional neural networks to correct artifacts in T2w images of the brain and achieved a reduction in mean squared error of 41.8% (26). However, to date, no studies have systematically trained for or evaluated the effects across a broad range of image corruption, and none have been explicitly developed for tumor molecular profile classifications.

The purpose of our study was twofold: 1) to evaluate the effect of motion corruption on deep-learning based molecular marker classification accuracy in brain gliomas, and 2) to determine if deep learning motion correction can recover classification accuracies to levels similar to the non-corrupted images. For motion correction, we developed a novel deep learning-based network to remove motion artifact from T2w brain MR images across a broad range of data corruption in glioma patients. In assessing the effects of motion artifact corruption on the classification accuracies, we utilized our previously developed deep learning networks for determination of IDH mutation status (7), 1p/19q codeletion (15), and MGMT methylation. These networks use only T2-weighted images and have provided the highest MRI-based classification accuracies reported to date, approaching those of invasive tissue-based histopathologic and molecular methods. The results of this study will elucidate the effects of motion artifact on deep learning-based molecular classification, and the relative importance of robust correction methods for facilitating clinical applicability. Figure 1 provides an overview of our study design. This consists of 1) Simulating motion in the original T2w glioma images 2) Training the network on the motion simulated images to generate artifact-free images using the non-distorted images as ground truth 3) Evaluating the performance of the network for correcting motion artifact on the held-out subjects and 4) Testing the performance of our previously developed glioma molecular classification networks using the motion-corrupted and motion-corrected images.

**Figure 1.**
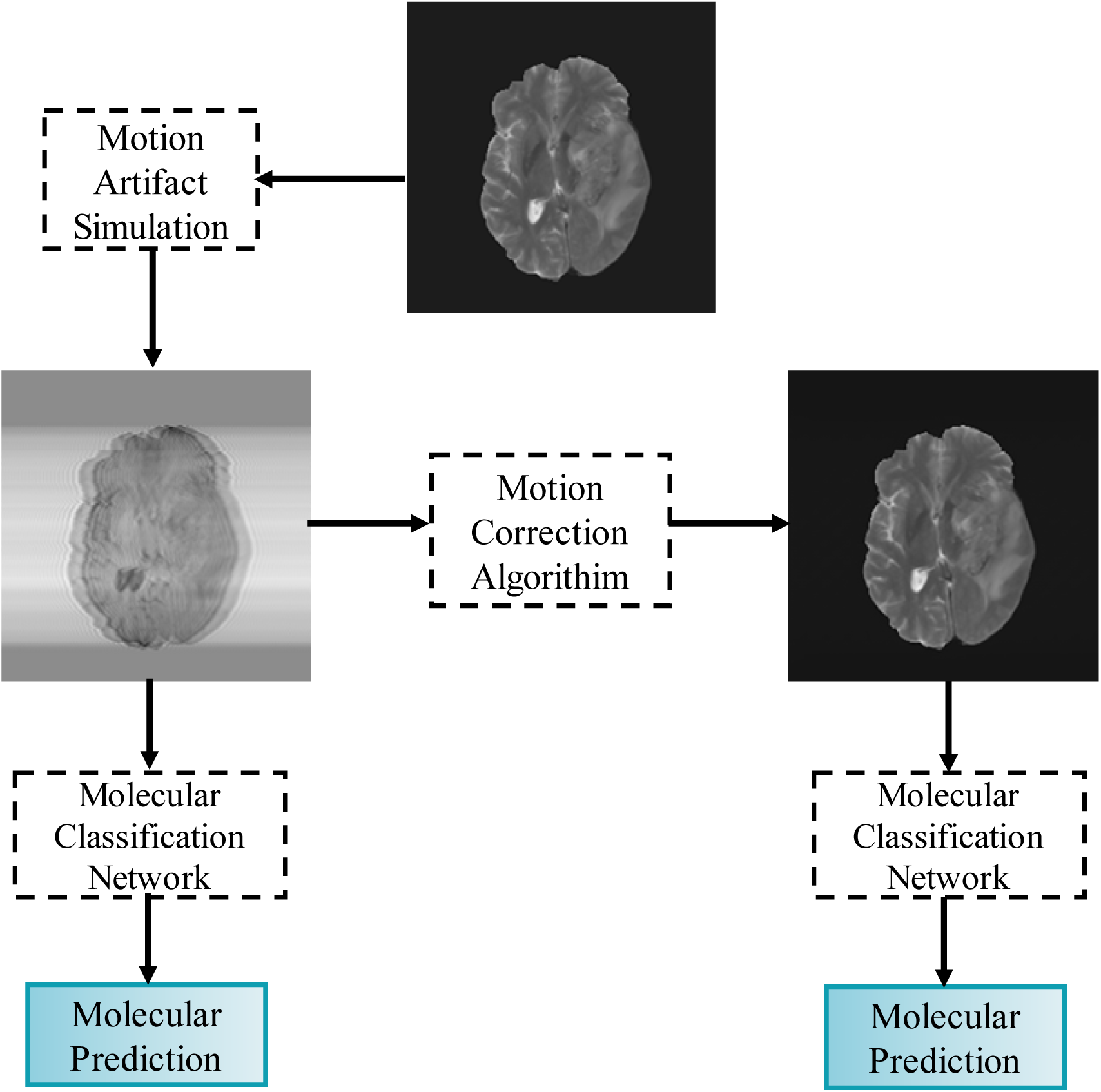
Study overview

## 2. Materials and Methods

### 2.1 Dataset and Preprocessing

Subject imaging data were retrieved from the TCIA database (27), while corresponding genomic information was obtained from the TCGA database (28). Only preoperative cases with T2w MR images were included in the study. The final IDH dataset consisted of 214 subjects (94 IDH-mutated and 120 IDH wild-type subjects). Imaging and genomic data from 368 subjects were obtained from the TCIA and TCGA databases with 1p/19q codeletion status (130 1p/19q co-deleted and 238 non-co-deleted) as well as 247 subjects with MGMT methylation status (163 MGMT methylated and 84 unmethylated). The TCGA subject IDs, molecular profile, age, and gender can be found in the Supplementary Data (Tables 1, 2, and 3).

**Table 1.**
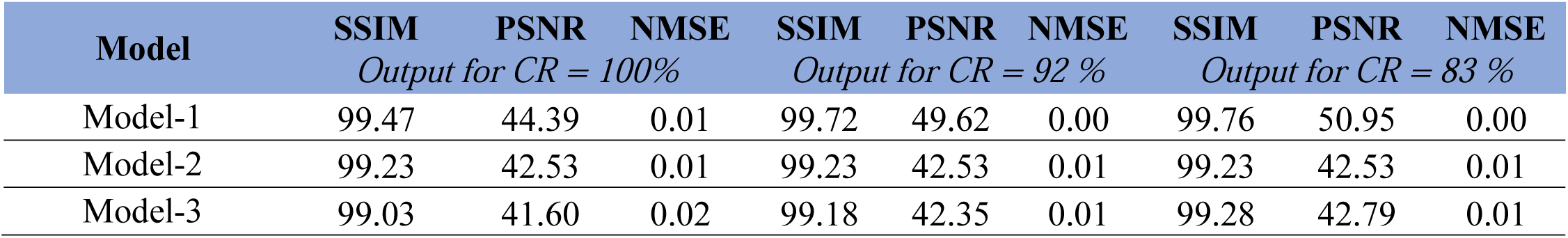
Motion correction model performance.

Minimal preprocessing was applied to the imaging datasets, consisting of (1) Co-registration of the T2w MR images to the SR124 T2w template (29) using Advance Normalization Tools (ANTS) software (30) (2) Skull stripping of the T2w images using the Brain Extraction Toolkit (BET; FMRIB software library [FSL]) (31-34) (3) N4 Bias Field Correction (35) to removing radiofrequency pulse inhomogeneity, and (4) Image intensity normalization to zero mean and unit variance (36). The preprocessing steps required less than 5 minutes per dataset.

### 2.2 Motion Simulation

Motion artifacts were simulated by adding additional phase to the k-space, which was obtained after applying an inverse Fourier transformation to the T2w image (37). The motion artifact incorporated into the k-space data closely simulates the additional phase induced by the patient movements encountered during MR imaging. Specifically, translational motion was incorporated into the image along the phase encoding direction using equation 1 below. Briefly, the k-space data along the phase encoding direction (k_y_) is multiplied with an exponential function to incorporate the additional phase (24) in the simulated k-space data. 

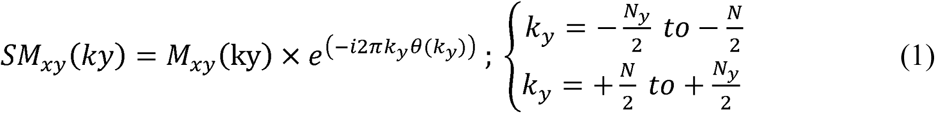

M_xy_(k_y_) is the original k-space, SM_xy_(k_y_) is the motion simulated k-space, and θ(k_y_) is the phase induced by motion. The total number of corrupted k-space lines is given by N, such that the outermost N/2 lines on either side of the k-space are corrupted. The corruption rate (CR) represents the percentage of corrupted lines in k-space (Fig. 2), where CR = N/Ny, with Ny being the total lines of phase encoding lines (e.g., N_y_ = 240). In our study, the number of corrupted k-space lines (N) ranged from 10, 20, 60, 80, 100, 120, 140, 150, 160, 180, 220, and 240, which corresponded to CRs of 4%, 8%, 25%, 33%, 42%, 50%, 58%, 63%, 67%, 75%, 83%, 92%, and 100%. These CR values were selected to represent a broad range of motion artifacts, from minimal to highly corrupted images. Additionally, in contrast to prior studies, all lines of k-spaces were utilized, including the low spatial frequencies contained in the central 7% of the k-space.

**Figure 2.**
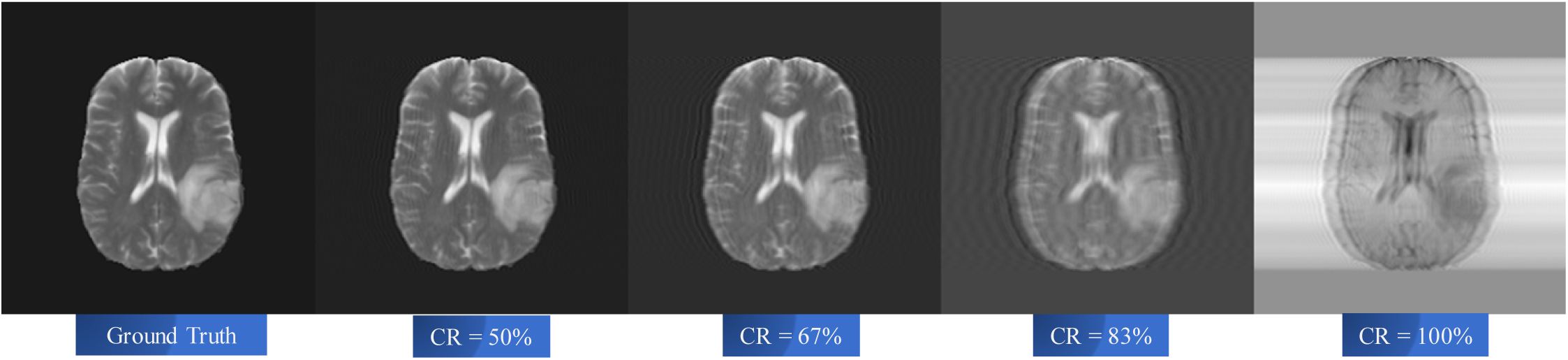
Example of simulated motion data. From left to right, ground truth T2w image (column 1) and corrupted images for CR=50%, 67%, 83% and 100% (columns 2 −5).

### 2.3 Network Architecture

#### Model-1 (Blur-Net)

Supplementary Figure 2 shows the network architecture for Model-1 (Blur-Net). The Model-1 motion correction algorithm is adapted from a 2D Dense-Unet architecture (38). It consists of 4 transition down blocks, and 4 transition up blocks with an initial and a final convolution layer. Each transition down block consists of a dense block and a pooling block, while each transition up block consists of an up-sampling block and dense block. Each dense block has 5 densely connected convolutional layers (39), where each layer is connected to every other layer in the dense block. The feature maps of all the convolutional layers in the dense block were concatenated to the output of the block, providing a dense connection. The output of the dense block was also concatenated with the input.

The encoder part of the network has 4 transition down blocks, which are comprised of 4 dense blocks and a subsequent pooling block. Each pooling block contains a batch normalization layer, activation layer, convolution layer, spatial drop out layer, and maximum pooling layer. The decoder part of the network has 4 transition up blocks, which are comprised of 4 dense blocks, each of which are preceded by an up-sampling block. Up-sampling blocks are comprised of a batch normalization layer, activation layer, deconvolution layer, and spatial dropout. The activation layer used was a rectified linear unit (ReLU). Tompson et al. (40) introduced the concept of spatial dropout, a regularization technique that drops random subsets of feature maps in the subsequent layer, for spatially co-related features in images. Dense block 1 (DB 1) was used as a bottleneck in the network, which helps reduce the number of feature maps. This reduction can assist with memory optimization, allowing the network to operate within the resource limits of the GPU. Additionally, as the number of features was reduced, the network can be trained in relatively less time. A total of 50 2D densely connected convolution layers were implemented in the Blur-Net architecture.

#### Model-2 (SE-Net 154)

The network architecture of Model-2 (Supplementary Figure 3) was based on a modified version of the squeeze and excitation network (SE-Net) (41). SE-Net has shown promising results in image classification, winning the ImageNet Large Scale Visual Recognition Challenge (ILSVRC) 2017 classification challenge (42). The SE-Net 154-based motion correction architecture consists of an input block, four transition down blocks (TD Block), and four transition up blocks. The input block consists of a 3-convolution layer, and a maximum pooling layer. Each transition down blocks consists of two convolution layers as well as a group convolution layer, concatenation layer, SE block, addition layer, and activation layer. The group convolution layer splits the input tensor into the number of groups, and then each group runs through the convolution layer. In our network, the input tensor was split into 64 groups. The final outputs of all the group convolutions are then concatenated. The SE block is comprised of a global average pooling layer, lambda layer (expanding the dimension), two convolution layers, ReLU activation layer, sigmoid activation layer, and a multiplication layer. A lambda layer was used for expanding the dimensions of the input tensor. Each transition up block consists of an up-sampling layer, concatenation layer, and two convolution layers. Each transition down block was iterated sequentially. TD Blocks 1, 2, 3, and 4 were iterated for 3, 8, 36, and 3, respectively. A total of 3407 2D convolution layers were implemented in the SE-Net 154 network architecture.

#### Model-3 (SE-Net 154) with Blur Loss

The network architecture for Model-3 is largely the same as Model-2, with the only difference being the use of a different loss function that incorporated the perceptive blur metric to evaluate image blurriness (43). The use of this metric was intended to help the algorithm learn and reduce errors in its predictions, particularly with regard to image sharpness.

### 2.4 Training

Training of the motion correction networks was performed in 2 phases. First, the three motion correction models were trained using 214 subjects from the IDH dataset. The subject data were randomly shuffled into three groups for training, in-training validation, and testing (∼71 subjects per set). Performance of all 3 networks was compared using the results on the held-out testing set. The best performing network was then selected and retrained using a larger combined dataset of 446 unique subjects from all three molecular marker groups (IDH, 1p/19q, and MGMT). This combined dataset was randomly shuffled into three groups to perform a 3-fold cross-validation. Supplementary Data (Table 4) shows the data distribution used for the combined cross-validation datasets. For each of the three folds, the groups were alternated between training, in-training validation, and held-out testing sets (∼149 subjects per set in the combined dataset). Motion correction network performance was reported based on the held-out testing set, which was not seen during the training step. Additionally, the 2D slices were separated by subject for each of the cross-validation folds. This latter procedure helps eliminate the problems of subject duplication and data leakage between the training and testing steps (44,45).

Data augmentation was performed on the input MR images to increase training quality and diversity, which helps for training models with limited data. The data augmentation steps included horizontal and vertical flipping of the images. The networks were implemented on NVIDIA Tesla V100s GPU and Keras (46), a python package with Tensorflow (47) as the backend, with an adaptive moment optimizer (Adam). The initial learning rate of the optimizer was set at 1 × 10^−5^. Model-1 and Model-2 were trained using a combined loss function of structural similarity index (SSIM) loss, peak signal-to-noise ratio (PSNR) loss, mean absolute error (MAE), and perceptual loss, with equal weighting for the structural components, noise level, and perceived image quality. The loss function for Model-3 differed in its use of blur loss instead of MAE and perceptual loss. All the networks were trained from scratch with a batch size of 4 on GPU V100s. Training time for all the networks was in the range of 96 to 120 hours.

### 2.5 Testing

The three trained motion correction networks were evaluated on the held-out testing set. Testing was carried out on Tesla V100 and P40 NVIDIA GPUs. The motion corrupted input image and predicted corrected output image were compared to the ground truth reference image. The performance of the models was evaluated using 1) SSIM which quantifies the perceived degradation in image quality between predicted and ground truth images by considering structural, luminance, and contrast features in the image 2) PSNR, an additional measure of image quality, where the signal is the ground truth image, and noise is the error in the predicted image that was created during image reconstruction by the network, and 3) normalized mean squared error (NMSE) which is used to evaluate the pixel-level difference between the predicted image and the ground truth image. The best performing motion correction network was then retrained on the combined dataset and evaluated on the held-out testing set for each of the three cross-validation folds, utilizing the same metrics as above. The results from each fold were averaged across all subjects for each corruption level. The testing time for each subject was less than 60 seconds.

Classification accuracy for IDH mutation status was initially performed for the three motion correction networks to further corroborate the selection of the best-performing network. Molecular classification accuracies for IDH, 1p/19q, and MGMT promoter were then evaluated using the retrained best-performing motion correction network. Accuracies were determined using the ground truth uncorrupted images and at each of the 13 image corruption levels (ranging from 4% CR – 100% CR) for each cross-validation fold using our previously trained deep learning T2w image-based molecular classification networks (7,15). The results were averaged across folds to provide a mean classification accuracy for each molecular marker at each corruption level. This process was then repeated on the motion-corrected images for IDH, 1p/19q, and MGMT for each corruption level to determine if the ground truth accuracies could be recovered.

## 3 Results

### 3.1 Comparison of motion correction algorithms

Table 1 shows the SSIM, PSNR, and NMSE metrics for our three motion correction networks on the held-out testing dataset at the three highest motion corruption levels. Figure 2 demonstrates the effects of several motion corruption levels on a T2w brain image. Model-1 (Blur-Net) achieved the best performance across all three metrics, most notably PSNR. Figure 3 shows the motion-corrected output images generated by the three networks for a single subject at high levels of motion corruption (corruption rate [CR]=83% and 100%). Although the three models generated similar results at low corruption levels (CR=50%), at higher corruption levels (CR =92%), Model-1 generated sharper images that were visually indistinguishable from the ground truth image (Figure 4 and Supplementary Figure 1). Model-1 surpassed the other two models in terms of quantitative metrics as well as perceived visual quality.

**Figure 3.**
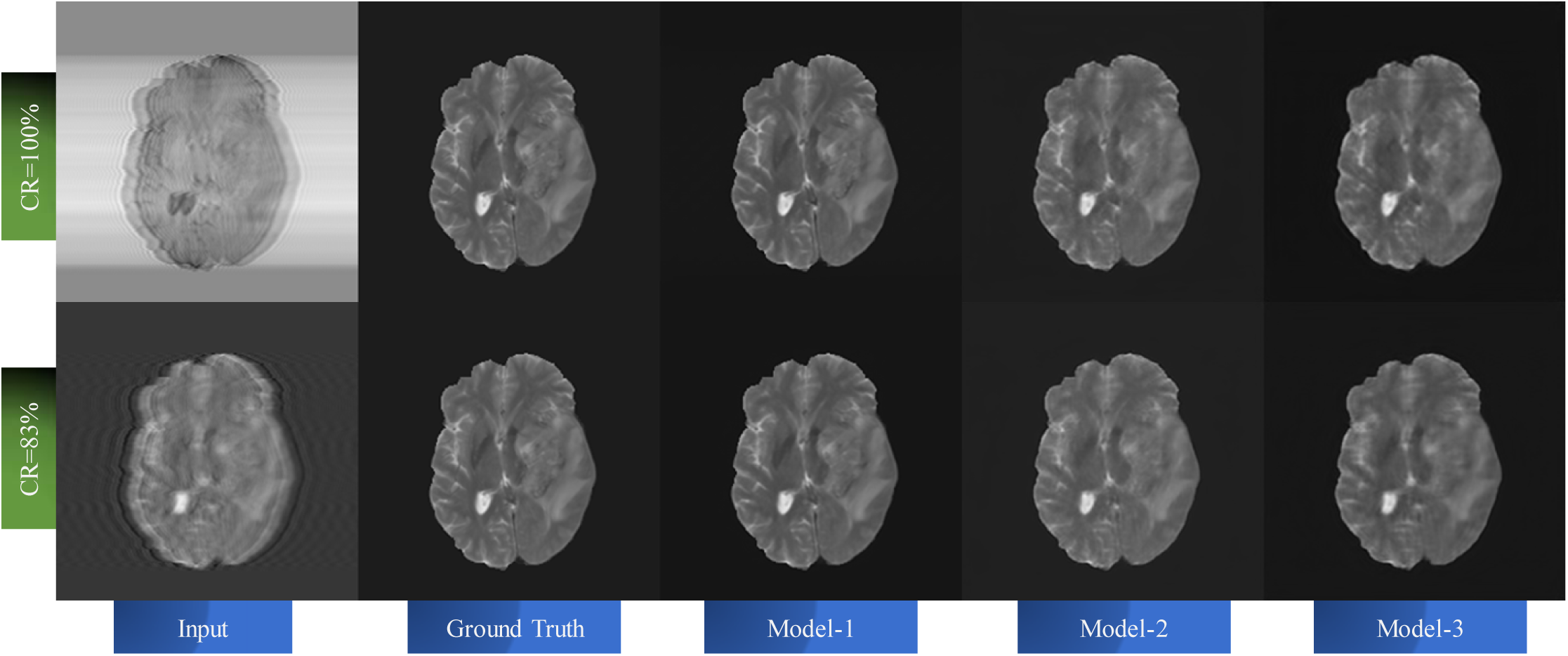
Examples of motion correction at high corruption levels for the three models. Input corrupted image (column 1) at CR of 83% (bottom row) and 100% (top row), ground truth (column 2), Model-1 output (column 3), Model-2 output (column 4), and Model-3 output (column 4). Model-1 provides visually obvious improved performance over the other two models.

**Figure 4.**
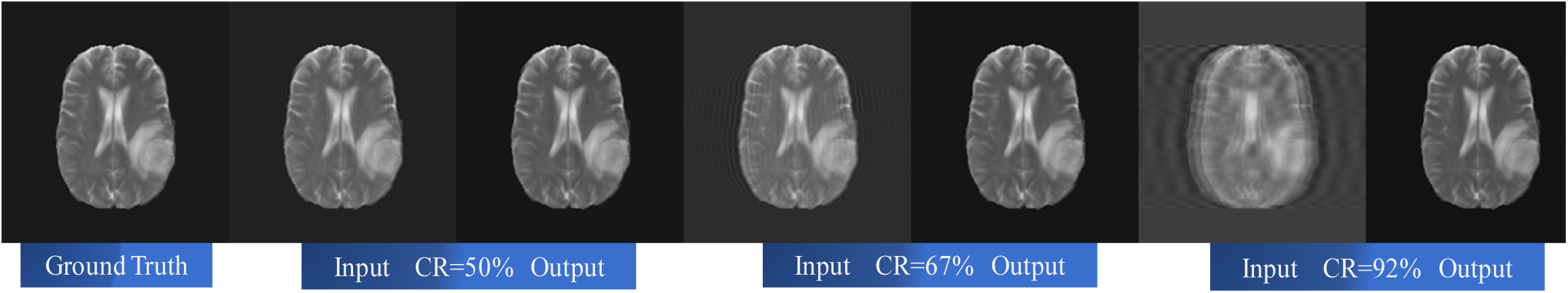
Model-1 motion correction performance for a single subject. From left to right, ground truth image (column 1), 50% CR input and corrected output (columns 2 and 3), 67% CR input and output (columns 4 and 5), 92% CR input and output (columns 6 and 7).

### 3.2 Effects of motion and motion correction on IDH, 1p/19q, and MGMT Classification Accuracy

Figure 5 compares IDH mutation status classification accuracies for the three motion correction networks using uncorrected motion corrupted images and motion-corrected images at increasing levels of motion corruption. The IDH classification begins to fail on the motion corrupted images at a CR of 42%, progressively decreasing thereafter. Similar to the analysis of quantitative metrics, Model-1 achieved the best results, maintaining a 97% IDH classification accuracy through a CR of 92%. While the other two models were able to improve classification accuracy over the full range of image corruption levels, they were unable to achieve the performance of Model-1.

**Figure 5.**
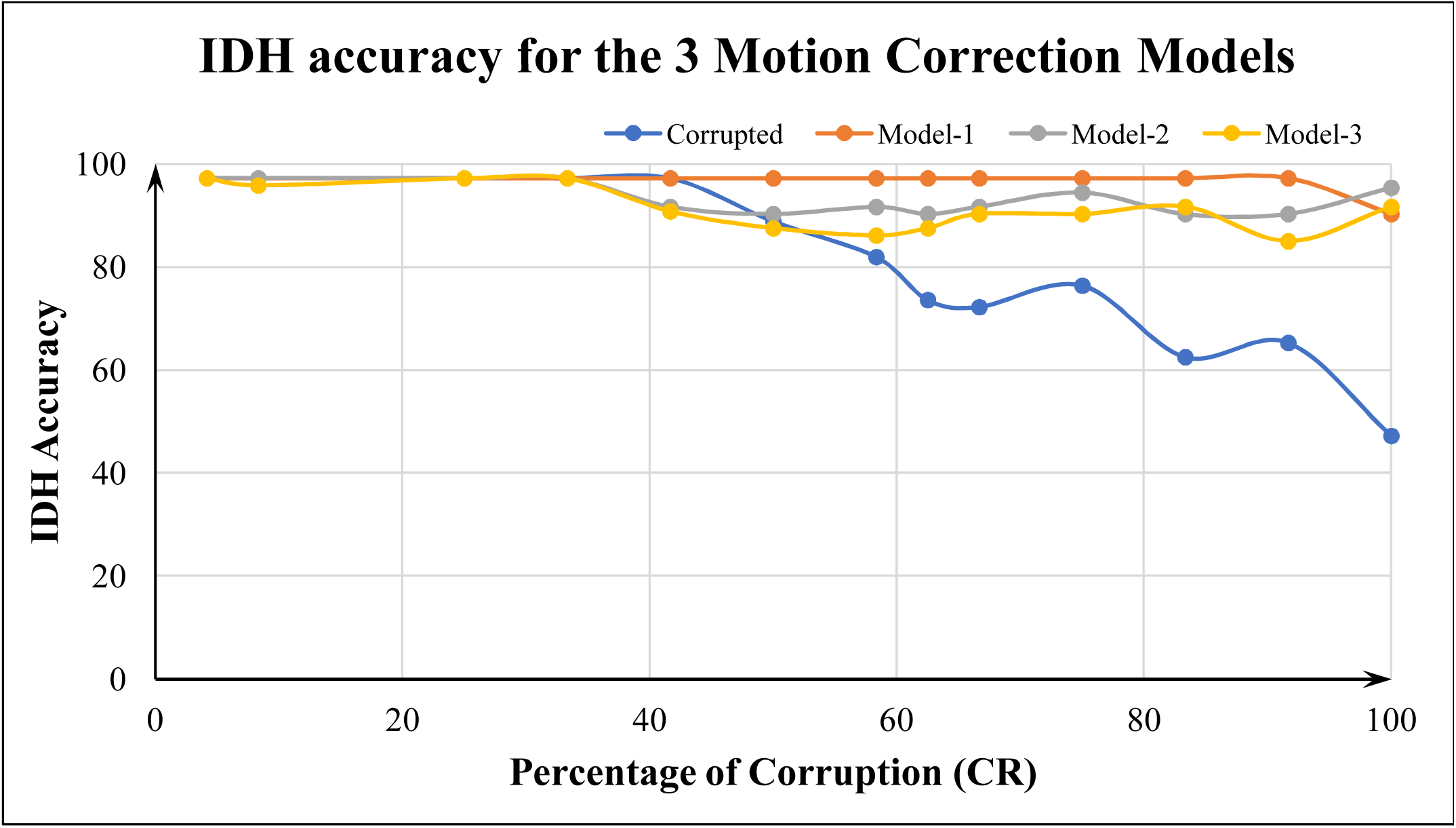
IDH classification accuracy and percent corruption for motion corrupted images and motion-corrected images for the three correction networks. Motion corrupted accuracies (blue), as well as accuracies following motion correction for Model-1(orange), Model-2 (grey), and Model-3 (yellow) are shown. A progressive decrease in classification accuracy for the corrupted images is demonstrated beyond 42% corruption (blue line). Model-1 performed best (orange line), recovering the original 97% classification accuracy out to 92% corruption level.

Figures 6 demonstrates the IDH, 1p/19q codeletion, and MGMT methylation status classification performance using the best performing network (Model-1) after it was retrained on the larger combined dataset. The classification accuracy on the corrupted images fell off at 42% corruption for both IDH and 1p/19q, while for MGMT, there is a sharp decline at 63% corruption. For the corrected images, IDH classification is maintained at 97% accuracy out to 92% corruption, and recovers to 94% accuracy even at 100% corruption. More remarkably, for correction of the native images and at lower levels of image corruption (0%-33%), IDH classification accuracy exceeded the performance of the uncorrupted images achieving up to 99% accuracy. For both 1p/19q and MGMT, 83% accuracy was recovered out to 92% corruption and 82% accuracy was recovered at 100% corruption.

**Figure 6.**
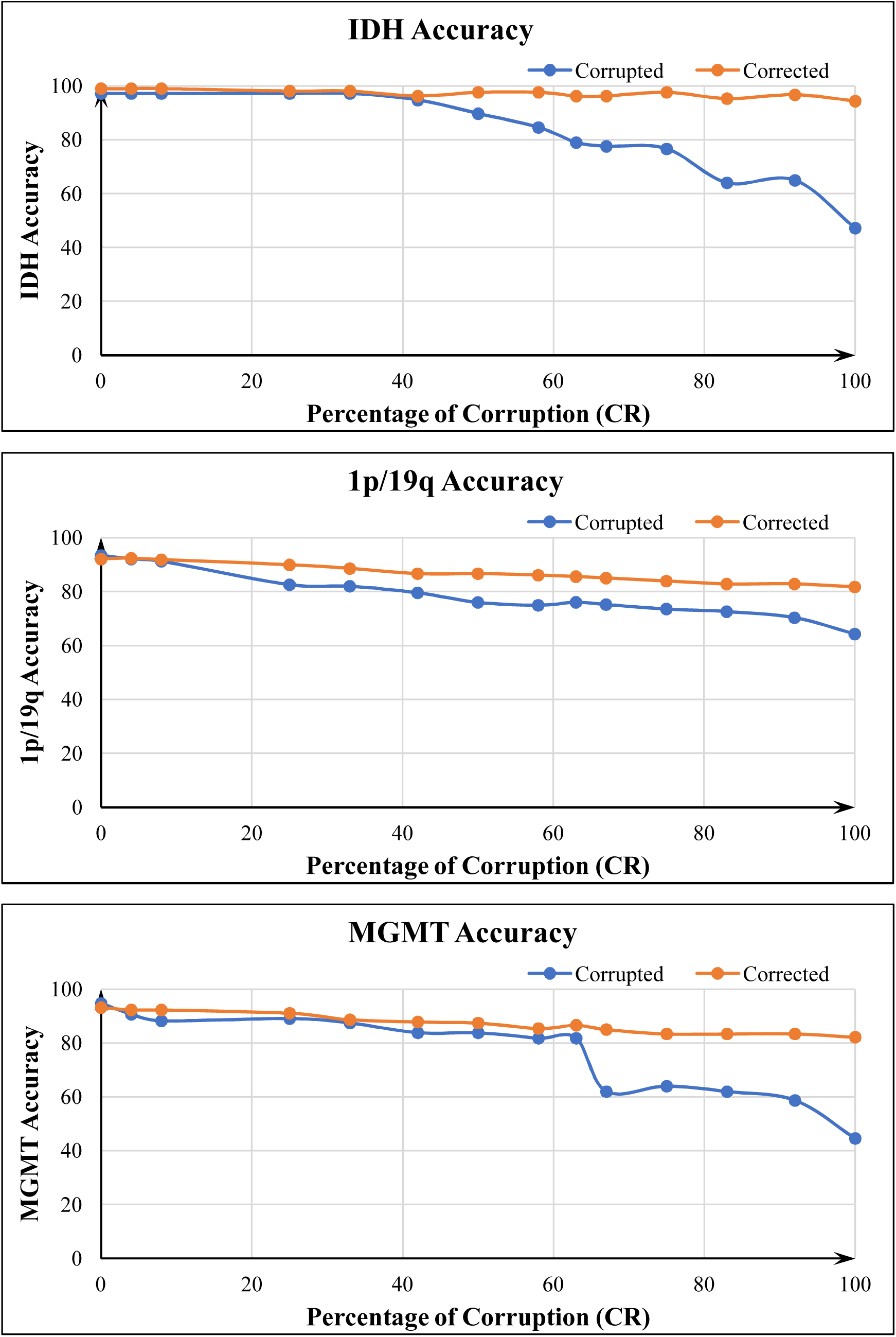
IDH, 1p/19q, and MGMT classification accuracies for uncorrected motion corrupted (blue lines) and Model-1 corrected images (orange lines) averaged across the 3-folds for each molecular marker. Recovery of accuracy was best for IDH classification, achieving 99% accuracy at low-levels of motion, and recovering the original 97% accuracy out to 92% corruption level.

In the case of IDH, we also investigated the change in voxel-wise Dice scores (Supplementary Data, Figure 4). The motion-corrected images demonstrated improved voxel-wise Dice scores across motion levels compared to the uncorrupted images. For the uncorrupted images, voxel-wise Dice scores were 0.86 and 0.87 for mutated and wild-type, respectively. Following correction, the IDH mutated Dice score increased to 0.88 at up to 25% corruption. Both IDH wild-type and IDH mutated dice scores were increased relative to the uncorrupted images even at 100% image corruption. Supplemental Figure 4 shows the voxel-wise Dice scores across different corruption levels for IDH mutated, IDH wild-type, and for the whole tumor segmentation (IDH mutated + IDH wild-type).

## 4 Discussion

We developed and tested three motion correction algorithms that were able to handle a broad range of motion corruption levels for T2w MR images of gliomas. We demonstrate that the performance of previously trained glioma molecular profile (IDH mutation, 1p/19q codeletion, and MGMT methylation) classifiers was adversely affected by motion corruption, with progressive loss in accuracy with increasing motion corruption. Importantly, classification accuracies could be recovered or significantly improved after applying the motion correction algorithm, even at very high levels of motion corruption. In the case of IDH classification, 99% accuracy was achieved following motion correction, exceeding the performance using ground truth images (which was 97%) (7).

Of the three motion correction algorithms compared, Model-1 (Blur-Net) performed the best. This model was based on a 2D Dense-Unet architecture, which may account for its superior performance. With its densely connected design, all feature maps are reused, such that each layer in the architecture received a direct supervision signal. In addition, the Dense-Net architecture alleviates the vanishing gradient problem in machine learning, which can prevent the neural network from further training. Other advantages include feature propagation and feature reuse, as described by Huang et al. (39).

Moreover, all three models achieved excellent performance with SSIM of over 0.99 and RMSE of less than 0.03 for all motion corruption levels. For comparison, Duffy et al. (24) achieved SSIM and RMSEs of 0.97 and 0.04, respectively, while Sommer et al. (26) reported SSIM of 0.86 to 0.924 when comparing motion-corrected brain MRIs to uncorrupted ground-truth images. Of note, Duffy et al. used brain masks to extract the brain-only regions from the MR Image prior to training. Masking of the image relies on preprocessing pipelines, which may not generate consistent or accurate outputs. Our approach involved correcting the entire image volume, including the brain, skull, and soft tissues, bypassing this preprocessing step and any potential errors associated with it.

In contrast to prior studies which used fixed or limited sets of motion corruption levels (Duffy et al. (24) limited corruption to 30 lines of k-space), we systematically trained our models using a broad range of motion corruption levels, from minimal (4%) to severely distorted with 100% of k-space lines affected. This approach better captures real-world conditions where there is a mixture of motion artifacts, from mild motion that largely preserves diagnostic information to more severe cases that are effectively uninterpretable. Our networks were able to accurately correct motion across these varying levels.

Another challenge facing the clinical application of deep learning networks is the generalizability of the algorithms. Although the TCIA dataset used to train the networks is relatively small compared to the size of databases typically utilized for deep learning applications, it nevertheless represents one of the largest publicly available brain tumor databases. The database also surpasses the size of the training datasets used in other brain MRI motion correction studies, which contributes to our overall improved performance. The issue of generalizability is further addressed as the TCIA cases were obtained from multiple institutions using different MRI vendor platforms with a variety of image acquisition parameters. The importance of generalizability is also highlighted by the improved results of the Blur-Net algorithm on IDH classification when trained on the combined dataset.

The algorithms we previously developed for brain tumor molecular classification were trained on relatively motion-artifact free images from the TCIA database. A baseline level of motion artifact may be present and, when averaged throughout the database, could partly account for the robustness of classification accuracy that was retained to a corruption level of CR=42%. However, the performance declined sharply at progressive image corruption levels. Although an additional approach would be to train using corrupted imaging data intentionally, this strategy could lead to the networks erroneously learning incorrect imaging features (in the form of motion corrupted imaging features or the motion artifacts themselves) as the basis for classifying the molecular markers. As such, deep learning image-based classification studies have excluded data with significant artifacts from their training database (6,48,49). Alternatively, conventional non-machine-learning based motion correction strategies could be utilized before applying the molecular classification algorithms. However, a key advantage to our deep learning approach is that it may be applied retrospectively to any previously acquired image without the need for any additional acquisition time, special scanner preparatory steps, or additional data input.

The TCIA dataset has a variety of gliomas with different biological behaviors, including glioblastoma multiforme (GBM), anaplastic astrocytoma, low-grade glioma, and oligodendroglioma, with their associated variations in IDH mutation, 1p/19q codeletion, and MGMT methylation status. Importantly, the networks, particularly Blur-Net, were able to not only remove the imaging artifacts but also preserve the key imaging features of the tumors from the MR images necessary for accurate classification. This was evidenced by the full recovery of classification accuracy for the IDH network extending out to a corruption level of 92%, and markedly improved accuracies for 1p/19q codeletion, and MGMT methylation networks following the application of the Blur-Net motion correction algorithm. For IDH classification, the motion-corrected images outperformed the ground truth images, achieving a remarkable accuracy of 99%. This may reflect the presence of latent image artifacts within some of the ground truth images, which were removed by the motion correction algorithm, facilitating classification performance. Additionally, there was an improvement in IDH mutant voxel-wise Dice scores following motion-correction compared to the uncorrupted native images, providing increased confidence for the subject-wise classification. These compelling results support the routine use of a deep learning-based image artifact removal step for imaging-based deep learning applications in the classification of glioma molecular profiles. We demonstrated that this implementation enhances the robustness of the classification pipeline to real-world challenges, which facilitates its potential clinical feasibility and implementation, without any loss of classification accuracy across corruption levels.

A significant additional benefit from the use of our motion correction algorithm is the improvement in image quality for interpretation by diagnostic radiologists *(50)*. As noted, the implementation of our algorithm is retrospective in nature and therefore does not require separate accommodations during the image acquisition step, which ultimately benefits the patient as well as the health care system by requiring less scan time. Furthermore, the robustness of the motion correction for an extensive range of artifact levels indicates that it can handle the patient heterogeneity encountered in clinical practice, significantly reducing the number of nondiagnostic scans, the scan time required that would have been required for prospective motion correction techniques, as well as the need for repeating scans. Altogether, these represent significant advancements from current practices, reducing the cost of clinical operation, improving clinical workflow, and ultimately improving patient care.

### Limitations and future directions

While we achieved excellent performance for recovering classification accuracy for glioma molecular profiles, our study was confined to the TCIA database. Before using our approach in the clinical environment, it will be essential to train and validate using additional independent datasets. The focus of our study was to specifically address the effect of motion artifacts in MR images on deep learning molecular classification, although we recognize that other artifacts such as magnetic field inhomogeneity, Gaussian noise, and radiofrequency spikes can also affect MR image quality. Adapting our approach to address these artifacts would not require significant modifications, and many of these can be simulated retrospectively on previously acquired imaging data for training.

Although our molecular classification networks performed better using motion-corrected images compared to motion-corrupted images, we were not able to fully restore the classification accuracies of the 1p/19q and MGMT networks achieved using uncorrupted images. These findings indicate that the three classification algorithms differed in terms of their resilience to motion artifact. Both the 1p/19q and MGMT networks were based on the trained IDH classifier network architecture, with fine-tuning to the decoder part of the network to adjust classification weightings without changes to the encoder part. This led to faster training and resultant excellent classification accuracies using uncorrupted images but appears to have also rendered the networks less robust to image corruption compared to the fully-trained IDH network. While we achieved superior motion correction results compared to previous studies, subtle residual artifacts within the image appear to have been sufficient to affect molecular classification performance. It is also possible that performance could be enhanced with modifications to the motion correction network architecture. We chose to use a 2D network design for the associated lower computational resource demands, as well as the fact that the TCIA database contained 2D T2w images. However, recent advances in deep learning network architecture, such as 3D architectures, could be adapted in the future.

## 5 Conclusion

We developed a deep learning-based motion correction algorithm (Blur-Net) using a state-of-the-art Dense-Unet architecture, which was trained and validated using T2w glioma images from the TCIA database. Blur-Net achieved superior performance correcting a broad range of motion-related image degradation, indicating the ability to handle the heterogeneity encountered in the clinical setting. We demonstrate that previously high-performing classification networks for IDH mutation status, 1p/19q codeletion, and MGMT methylation progressively lose accuracy with increasing motion-related image degradation. However, by incorporating Blur-Net motion correction prior to the classification step, full recovery of classification accuracy was possible even at the highest degrees of motion disruption, indicating that not only was the network successful at removing artifacts but also in recovering crucial imaging features of the tumors. After training the motion correction network on a larger dataset composed of all three glioma markers, we achieved 99% accuracy for IDH classification, representing a new benchmark in non-invasive image-based IDH classification performance. Additionally, by correcting motion artifacts that would otherwise hinder interpretation by diagnostic radiologists, our algorithm may improve clinical workflow by reducing the number of nondiagnostic exams and the need for expensive repeat imaging. Blur-Net can be retrospectively applied to any suitable MR image and does not require additional scan times or special accommodations, which further facilitates potential clinical implementation.

