## Supplementary Figure 1, Figure 2, Figure 3, Figure 4, Supplementary Data (Tables 1, 2, and 3), Supplementary Data (Table 4) for "Brain Tumor IDH, 1p/19q, and MGMT Molecular Classification Using MRI-based Deep Learning: Effect of Motion and Motion Correction"

**SUPPLEMENTAL MATERIAL**

**FIGURES**

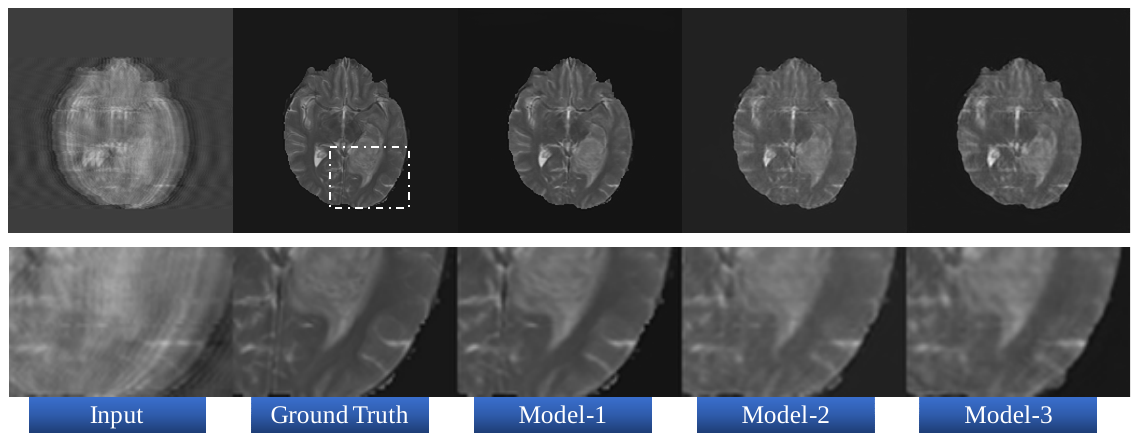

Figure 1. Example motion correction at high corruption level (CR = 92%) for the 3 models. Input corrupted image (column one), ground truth (column 2), Model-1 output (column 3), Model-2 output (column 4), and Model-3 output (column 5). Model-1 provided visually obvious improved performance over the other 2 models as evidenced by sharpness of the sulci and tumor borders.

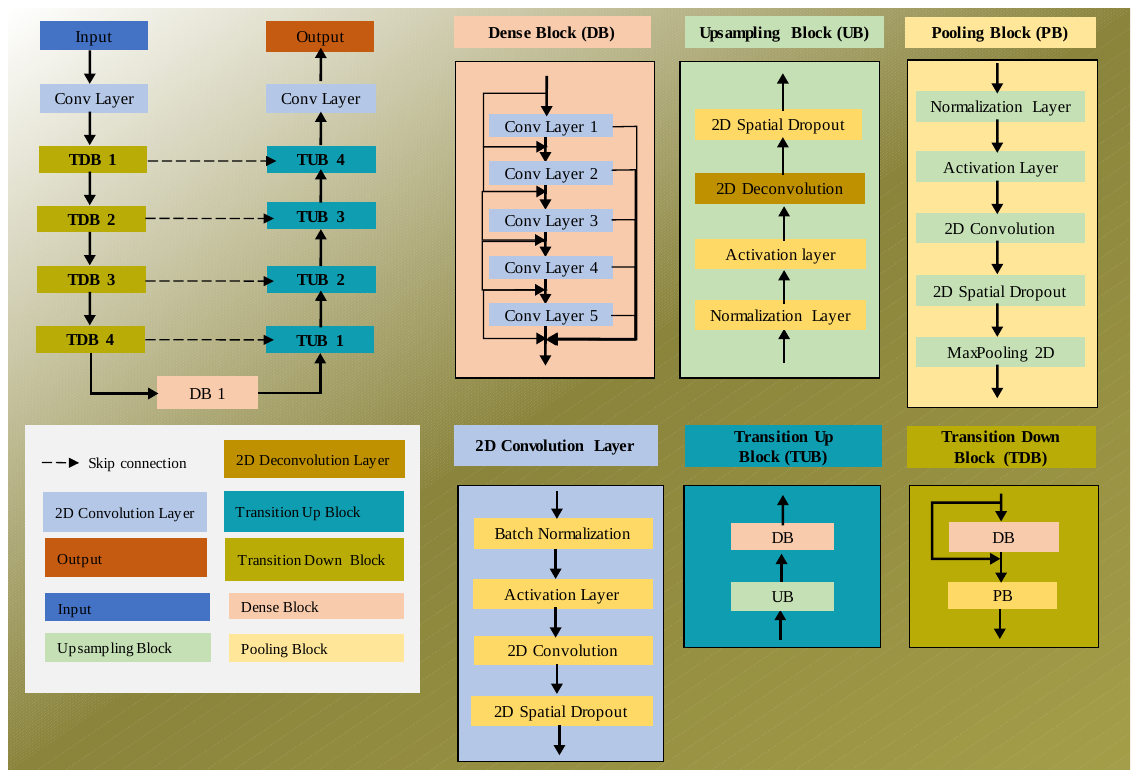

Figure 2. Architecture of the Blur-Net network (Model-1).

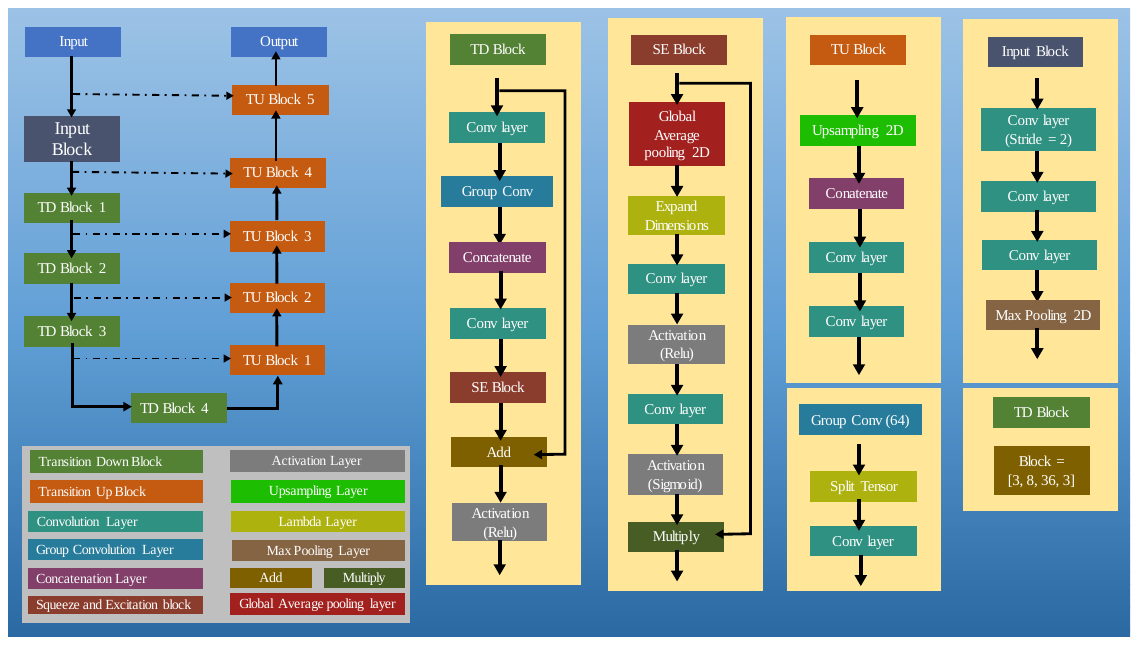

Figure 3. Architecture of SE-Net 154 (Model-2 and Model-3). For Model-3, the perceptive blur metric (blur loss) was substituted as the loss function.

Figure 4. IDH mutated, IDH wildtype, and whole tumor voxel-wise Dice scores for uncorrected motion corrupted (blue lines) and Model-1 motion corrected images (orange lines). IDH mutated and IDH wildtype Dice scores improve following motion correction.

IDH Mutated Dice Scores

Dice Score

| Percentage of Corruption (CR) | Corrupted | Corrected |
| --- | --- | --- |
| 0 | 0.864535 | 0.882767 |
| 4 | 0.851522 | 0.881916 |
| 8 | 0.868698 | 0.881164 |
| 25 | 0.850414 | 0.875384 |
| 33 | 0.850956 | 0.866767 |
| 42 | 0.825861 | 0.852333 |
| 50 | 0.762583 | 0.84883 |
| 58 | 0.693758 | 0.849568 |
| 63 | 0.659526 | 0.847497 |
| 67 | 0.63966 | 0.843434 |
| 75 | 0.602884 | 0.836209 |
| 83 | 0.509527 | 0.820737 |
| 92 | 0.448664 | 0.826882 |
| 100 | 0.277893 | 0.831307 |

IDH Wildtype Dice Scores

Dice Score

| Percentage of Corruption (CR) | Corrupted | Corrected |
| --- | --- | --- |
| 0 | 0.867025 | 0.8614 |
| 4 | 0.84774 | 0.854109 |
| 8 | 0.831026 | 0.853924 |
| 25 | 0.856045 | 0.843919 |
| 33 | 0.84201 | 0.83402 |
| 42 | 0.811117 | 0.827223 |
| 50 | 0.780255 | 0.818677 |
| 58 | 0.741334 | 0.811543 |
| 63 | 0.718896 | 0.801986 |
| 67 | 0.684783 | 0.806458 |
| 75 | 0.566014 | 0.806789 |
| 83 | 0.435969 | 0.798571 |
| 92 | 0.393876 | 0.799617 |
| 100 | 0.239077 | 0.777802 |

Whole Tumor Dice Scores

Dice Score

| Percentage of Corruption (CR) | Corrupted | Corrected |
| --- | --- | --- |
| 0 | 0.85 | 0.8691 |
| 4 | 0.866988 | 0.866403 |
| 8 | 0.860917 | 0.865739 |
| 25 | 0.862058 | 0.860172 |
| 33 | 0.851514 | 0.854012 |
| 42 | 0.832051 | 0.846985 |
| 50 | 0.804309 | 0.842092 |
| 58 | 0.765776 | 0.836819 |
| 63 | 0.741827 | 0.833184 |
| 67 | 0.703489 | 0.835123 |
| 75 | 0.595033 | 0.832134 |
| 83 | 0.44128 | 0.82338 |
| 92 | 0.409638 | 0.826829 |
| 100 | 0.368332 | 0.820314 |

**TABLES**

Table 1: Subject wise IDH mutation status and clinical variables

| **SUBJECT ID** | **Age** | **Gender** | **Histology** | **Grade** | **IDH**  **status** | **IDH**  **Allele** | **1p/19q co-deletion** | **Survival (months)** | **Karnofsky**  **Performance score** | **Cross-validation group** |
| --- | --- | --- | --- | --- | --- | --- | --- | --- | --- | --- |
| TCGA-02-0003 | 50 | male | glioblastoma | G4 | WT | N/A | non-codel | 4.7311 | 100 | 1 |
| TCGA-02-0006 | 56 | female | glioblastoma | G4 | WT | N/A | non-codel | 18.333 | 80 | 3 |
| TCGA-02-0009 | 61 | female | glioblastoma | G4 | WT | N/A | non-codel | 10.5793 | 80 | 1 |
| TCGA-02-0011 | 18 | female | glioblastoma | G4 | WT | N/A | non-codel | 20.6986 | 80 | 1 |
| TCGA-02-0027 | 33 | female | glioblastoma | G4 | WT | N/A | non-codel | 12.1563 | 100 | 2 |
| TCGA-02-0033 | 54 | male | glioblastoma | G4 | WT | N/A | non-codel | 2.8255 | 100 | 1 |
| TCGA-02-0034 | 60 | male | glioblastoma | G4 | WT | N/A | non-codel | 14.1276 | 80 | 3 |
| TCGA-02-0037 | 74 | female | glioblastoma | G4 | WT | N/A | non-codel | 3.614 | 80 | 1 |
| TCGA-02-0046 | 61 | male | glioblastoma | G4 | WT | N/A | non-codel | 6.8667 | 60 | 1 |
| TCGA-02-0047 | 78 | male | glioblastoma | G4 | WT | N/A | non-codel | 14.719 | 80 | 1 |
| TCGA-02-0048 | 80 | male | glioblastoma | G4 | WT | N/A | non-codel | 3.2198 | NaN | 2 |
| TCGA-02-0054 | 44 | female | glioblastoma | G4 | WT | N/A | non-codel | 6.5381 | 80 | 3 |
| TCGA-02-0060 | 66 | female | glioblastoma | G4 | WT | N/A | non-codel | 6.0124 | 80 | 1 |
| TCGA-02-0064 | 50 | male | glioblastoma | G4 | WT | N/A | non-codel | 19.7129 | 100 | 3 |
| TCGA-02-0068 | 57 | male | glioblastoma | G4 | WT | N/A | non-codel | 26.4153 | 80 | 1 |
| TCGA-02-0069 | 31 | female | glioblastoma | G4 | WT | N/A | non-codel | 28.6823 | 80 | 2 |
| TCGA-02-0070 | 70 | male | glioblastoma | G4 | WT | N/A | non-codel | 25.0354 | 80 | 2 |
| TCGA-02-0075 | 63 | male | glioblastoma | G4 | WT | N/A | non-codel | 20.83 | 80 | 2 |
| TCGA-02-0085 | 63 | female | glioblastoma | G4 | WT | N/A | non-codel | 51.2865 | 80 | 3 |
| TCGA-02-0086 | 45 | female | glioblastoma | G4 | WT | N/A | non-codel | 8.8051 | 100 | 1 |
| TCGA-02-0102 | 42 | male | glioblastoma | G4 | WT | N/A | non-codel | 27.0067 | 100 | 1 |
| TCGA-06-0119 | 81 | female | glioblastoma | G4 | WT | N/A | non-codel | 2.6941 | NaN | 2 |
| TCGA-06-0122 | 84 | female | glioblastoma | G4 | WT | N/A | non-codel | 6.1439 | NaN | 3 |
| TCGA-06-0127 | 67 | male | glioblastoma | G4 | WT | N/A | non-codel | 3.9754 | 60 | 1 |
| TCGA-06-0128 | 66 | male | glioblastoma | G4 | Mutant | IDH1 | non-codel | 22.7027 | 80 | 1 |
| TCGA-06-0129 | 30 | male | glioblastoma | G4 | Mutant | IDH1 | non-codel | 33.6434 | 100 | 1 |
| TCGA-06-0130 | 54 | male | glioblastoma | G4 | WT | N/A | non-codel | 12.9448 | 80 | 3 |
| TCGA-06-0132 | 49 | male | glioblastoma | G4 | WT | N/A | non-codel | 25.3311 | NaN | 3 |
| TCGA-06-0133 | 64 | male | glioblastoma | G4 | WT | N/A | non-codel | 14.2919 | NaN | 1 |
| TCGA-06-0137 | 63 | female | glioblastoma | G4 | WT | N/A | non-codel | 26.6782 | NaN | 3 |
| TCGA-06-0138 | 43 | male | glioblastoma | G4 | WT | N/A | non-codel | 24.2141 | 80 | 1 |
| TCGA-06-0139 | 40 | male | glioblastoma | G4 | WT | N/A | non-codel | 11.8935 | 60 | 1 |
| TCGA-06-0142 | 81 | male | glioblastoma | G4 | WT | N/A | non-codel | 2.2013 | NaN | 2 |
| TCGA-06-0143 | 58 | male | glioblastoma | G4 | WT | N/A | non-codel | 11.7292 | 60 | 2 |
| TCGA-06-0145 | 53 | female | glioblastoma | G4 | WT | N/A | non-codel | 2.3327 | NaN | 3 |
| TCGA-06-0147 | 51 | female | glioblastoma | G4 | WT | N/A | non-codel | 17.7745 | NaN | 2 |
| TCGA-06-0154 | 54 | male | glioblastoma | G4 | WT | N/A | non-codel | 13.9305 | 100 | 2 |
| TCGA-06-0157 | 63 | female | glioblastoma | G4 | WT | N/A | non-codel | 3.1869 | 40 | 2 |
| TCGA-06-0158 | 73 | male | glioblastoma | G4 | WT | N/A | non-codel | 10.8093 | 80 | 1 |
| TCGA-06-0166 | 51 | male | glioblastoma | G4 | WT | N/A | non-codel | 5.8482 | NaN | 1 |
| TCGA-06-0168 | 59 | female | glioblastoma | G4 | WT | N/A | non-codel | 19.6472 | 100 | 2 |
| TCGA-06-0174 | 54 | male | glioblastoma | G4 | WT | N/A | non-codel | 3.2198 | 80 | 2 |
| TCGA-06-0176 | 34 | male | glioblastoma | G4 | WT | N/A | non-codel | 51.3194 | 80 | 2 |
| TCGA-06-0184 | 63 | male | glioblastoma | G4 | WT | N/A | non-codel | 40.3458 | 80 | 2 |
| TCGA-06-0185 | 54 | male | glioblastoma | G4 | WT | N/A | non-codel | 36.9946 | 100 | 3 |
| TCGA-06-0187 | 69 | male | glioblastoma | G4 | WT | N/A | non-codel | 27.2039 | 60 | 3 |
| TCGA-06-0188 | 71 | male | glioblastoma | G4 | WT | N/A | non-codel | 28.4523 | 100 | 1 |
| TCGA-06-0189 | 55 | male | glioblastoma | G4 | WT | N/A | non-codel | 15.4089 | NaN | 1 |
| TCGA-06-0190 | 62 | male | glioblastoma | G4 | WT | N/A | non-codel | 10.415 | 80 | 2 |
| TCGA-06-0192 | 58 | male | glioblastoma | G4 | WT | N/A | non-codel | 18.3002 | 100 | 3 |
| TCGA-06-0213 | 55 | female | glioblastoma | G4 | WT | N/A | non-codel | 0.52568 | NaN | 3 |
| TCGA-06-0237 | 75 | female | glioblastoma | G4 | WT | N/A | non-codel | 13.6348 | NaN | 1 |
| TCGA-06-0238 | 46 | male | glioblastoma | G4 | WT | N/A | non-codel | 13.3062 | 80 | 2 |
| TCGA-06-0241 | 65 | female | glioblastoma | G4 | WT | N/A | non-codel | 14.949 | 100 | 1 |
| TCGA-06-0644 | 71 | male | glioblastoma | G4 | WT | N/A | non-codel | 12.3206 | 80 | 1 |
| TCGA-06-0645 | 55 | female | glioblastoma | G4 | WT | N/A | non-codel | 5.7496 | NaN | 3 |
| TCGA-06-0646 | 60 | male | glioblastoma | G4 | WT | N/A | non-codel | 5.7496 | 80 | 2 |
| TCGA-06-0648 | 77 | male | glioblastoma | G4 | WT | N/A | non-codel | 9.7908 | 80 | 3 |
| TCGA-06-0649 | 73 | female | glioblastoma | G4 | WT | N/A | non-codel | 2.1027 | NaN | 3 |
| TCGA-06-1806 | 47 | male | glioblastoma | G4 | WT | N/A | non-codel | 15.3104 | 90 | 2 |
| TCGA-06-2570 | 21 | female | glioblastoma | G4 | Mutant | IDH1 | non-codel | 9.3636 | 100 | 1 |
| TCGA-06-5408 | 54 | female | glioblastoma | G4 | WT | N/A | non-codel | 11.7292 | 80 | 3 |
| TCGA-06-5412 | 78 | female | glioblastoma | G4 | WT | N/A | non-codel | 4.534 | 80 | 2 |
| TCGA-06-5413 | 67 | male | glioblastoma | G4 | WT | N/A | non-codel | 8.8051 | 60 | 3 |
| TCGA-06-5417 | 45 | female | glioblastoma | G4 | Mutant | IDH1 | NA | 5.0925 | 80 | 2 |
| TCGA-06-6389 | 49 | female | glioblastoma | G4 | Mutant | IDH1 | non-codel | 7.7866 | 100 | 2 |
| TCGA-08-0390 | 69 | male | glioblastoma | G4 | WT | N/A | non-codel | 13.9633 | 60 | 3 |
| TCGA-12-0616 | 36 | female | glioblastoma | G4 | WT | N/A | non-codel | 14.719 | 100 | 2 |
| TCGA-12-0829 | 75 | male | glioblastoma | G4 | WT | N/A | non-codel | 20.5672 | 80 | 2 |
| TCGA-12-1093 | 66 | female | glioblastoma | G4 | WT | N/A | non-codel | 15.9675 | 80 | 3 |
| TCGA-12-1598 | 75 | female | glioblastoma | G4 | WT | N/A | non-codel | 15.6389 | NaN | 2 |
| TCGA-12-1601 | NaN | NA | glioblastoma | NA | WT | N/A | NA | NaN | NaN | 1 |
| TCGA-12-1602 | 58 | male | glioblastoma | G4 | WT | N/A | non-codel | 6.7681 | 60 | 1 |
| TCGA-12-3650 | 46 | male | glioblastoma | G4 | WT | N/A | non-codel | 10.9407 | 80 | 1 |
| TCGA-14-0789 | 54 | male | glioblastoma | G4 | WT | N/A | non-codel | 11.2364 | 40 | 3 |
| TCGA-14-1456 | 23 | male | glioblastoma | G4 | Mutant | IDH1 | non-codel | 40.9372 | 80 | 2 |
| TCGA-14-1794 | 59 | male | glioblastoma | G4 | WT | N/A | non-codel | 0.98565 | NaN | 3 |
| TCGA-14-1829 | 57 | male | glioblastoma | G4 | WT | N/A | non-codel | 7.1624 | 60 | 2 |
| TCGA-14-3477 | 38 | female | glioblastoma | G4 | WT | N/A | non-codel | 3.7783 | 80 | 1 |
| TCGA-19-1388 | 58 | male | glioblastoma | G4 | WT | N/A | non-codel | 12.9448 | NaN | 1 |
| TCGA-19-1390 | 63 | female | glioblastoma | G4 | WT | N/A | non-codel | 25.364 | 60 | 1 |
| TCGA-19-1789 | 69 | female | glioblastoma | G4 | WT | N/A | non-codel | 3.2526 | 60 | 2 |
| TCGA-19-2624 | 51 | male | glioblastoma | G4 | WT | N/A | non-codel | 0.16427 | NaN | 3 |
| TCGA-19-2631 | 74 | female | glioblastoma | G4 | WT | N/A | non-codel | 6.9981 | 60 | 2 |
| TCGA-19-5954 | 72 | female | glioblastoma | G4 | WT | N/A | non-codel | 7.9509 | 60 | 3 |
| TCGA-19-5958 | 56 | male | glioblastoma | G4 | WT | N/A | non-codel | 5.3882 | 80 | 2 |
| TCGA-27-1835 | 53 | female | glioblastoma | G4 | WT | N/A | non-codel | 21.29 | 80 | 2 |
| TCGA-27-1838 | 59 | female | glioblastoma | G4 | WT | N/A | non-codel | 11.4992 | 80 | 2 |
| TCGA-76-4926 | 68 | male | glioblastoma | G4 | WT | N/A | non-codel | 4.534 | 80 | 3 |
| TCGA-76-4932 | 50 | female | glioblastoma | G4 | WT | N/A | NA | 47.9024 | 80 | 2 |
| TCGA-76-4934 | 66 | female | glioblastoma | G4 | WT | N/A | non-codel | 2.5298 | 80 | 3 |
| TCGA-76-4935 | 52 | female | glioblastoma | G4 | WT | N/A | non-codel | 10.7764 | 80 | 3 |
| TCGA-76-6191 | 57 | male | glioblastoma | G4 | WT | N/A | non-codel | 16.6903 | 80 | 1 |
| TCGA-76-6192 | 74 | male | glioblastoma | G4 | WT | N/A | non-codel | 3.2855 | 80 | 1 |
| TCGA-76-6193 | 78 | male | glioblastoma | G4 | WT | N/A | non-codel | 2.6941 | 60 | 1 |
| TCGA-76-6280 | 57 | male | glioblastoma | G4 | WT | N/A | non-codel | 11.3678 | 80 | 2 |
| TCGA-76-6282 | 63 | male | glioblastoma | G4 | WT | N/A | non-codel | 17.0517 | 80 | 2 |
| TCGA-76-6285 | 64 | female | glioblastoma | G4 | WT | N/A | non-codel | 8.3451 | 80 | 2 |
| TCGA-76-6656 | 66 | male | glioblastoma | G4 | WT | N/A | non-codel | 4.8297 | 60 | 3 |
| TCGA-76-6657 | 74 | male | glioblastoma | G4 | WT | N/A | non-codel | 5.0268 | 80 | 1 |
| TCGA-76-6661 | 54 | male | glioblastoma | G4 | WT | N/A | non-codel | 0.22998 | 60 | 3 |
| TCGA-76-6662 | 58 | male | glioblastoma | G4 | WT | N/A | non-codel | 9.2651 | 80 | 1 |
| TCGA-76-6663 | 44 | female | glioblastoma | G4 | WT | N/A | non-codel | 7.7209 | 80 | 1 |
| TCGA-76-6664 | 49 | female | glioblastoma | G4 | WT | N/A | non-codel | 7.7866 | 80 | 1 |
| TCGA-CS-4941 | 67 | male | astrocytoma | G3 | WT | N/A | non-codel | 7.688 | 90 | 3 |
| TCGA-CS-4942 | 44 | female | astrocytoma | G3 | Mutant | IDH1 | non-codel | 43.8613 | 90 | 2 |
| TCGA-CS-4943 | 37 | male | astrocytoma | G3 | Mutant | IDH1 | non-codel | 18.1359 | 50 | 3 |
| TCGA-CS-4944 | 50 | male | astrocytoma | G2 | Mutant | IDH1 | non-codel | 10.6121 | 90 | 1 |
| TCGA-CS-5393 | 39 | male | astrocytoma | G3 | Mutant | IDH1 | non-codel | 40.1487 | 100 | 2 |
| TCGA-CS-5395 | 43 | male | oligodendroglioma | G2 | WT | N/A | non-codel | 20.9943 | 90 | 3 |
| TCGA-CS-5396 | 53 | female | oligodendroglioma | G3 | Mutant | IDH1 | codel | 9.955 | 90 | 2 |
| TCGA-CS-5397 | 54 | female | astrocytoma | G3 | WT | N/A | non-codel | 6.3739 | 80 | 2 |
| TCGA-CS-6186 | 58 | male | oligoastrocytoma | G3 | WT | N/A | non-codel | 17.6759 | 90 | 3 |
| TCGA-CS-6188 | 48 | male | astrocytoma | G3 | WT | N/A | non-codel | 23.8198 | 90 | 3 |
| TCGA-CS-6290 | 31 | male | astrocytoma | G3 | Mutant | IDH1 | non-codel | 17.9388 | 90 | 2 |
| TCGA-CS-6665 | 51 | female | astrocytoma | G3 | Mutant | IDH1 | non-codel | 12.4192 | 90 | 3 |
| TCGA-CS-6666 | 22 | male | astrocytoma | G3 | Mutant | IDH1 | non-codel | 8.4766 | 90 | 1 |
| TCGA-CS-6667 | 39 | female | astrocytoma | G2 | Mutant | IDH1 | non-codel | 7.5566 | 90 | 1 |
| TCGA-CS-6668 | 57 | female | oligodendroglioma | G2 | Mutant | IDH1 | codel | 8.0166 | 90 | 1 |
| TCGA-CS-6669 | 26 | female | oligodendroglioma | G2 | WT | N/A | non-codel | 7.3924 | 90 | 1 |
| TCGA-DU-5849 | 48 | male | oligodendroglioma | G2 | Mutant | IDH1 | codel | 14.5547 | NaN | 2 |
| TCGA-DU-5851 | 40 | female | oligoastrocytoma | G3 | Mutant | IDH1 | non-codel | 17.446 | 90 | 3 |
| TCGA-DU-5852 | 61 | female | oligoastrocytoma | G3 | WT | N/A | non-codel | 6.7353 | 80 | 2 |
| TCGA-DU-5853 | 29 | male | oligoastrocytoma | G2 | Mutant | IDH1 | non-codel | 13.3719 | 100 | 1 |
| TCGA-DU-5854 | 57 | female | astrocytoma | G3 | WT | N/A | non-codel | 8.4437 | 90 | 1 |
| TCGA-DU-5855 | 49 | female | oligoastrocytoma | G3 | Mutant | IDH1 | non-codel | 6.801 | 100 | 2 |
| TCGA-DU-5871 | 37 | female | oligoastrocytoma | G2 | Mutant | IDH1 | non-codel | 18.9244 | 100 | 3 |
| TCGA-DU-5872 | 43 | female | oligoastrocytoma | G2 | Mutant | IDH1 | non-codel | 17.4788 | NaN | 1 |
| TCGA-DU-5874 | 62 | female | oligodendroglioma | G2 | Mutant | IDH1 | codel | 15.1461 | 100 | 1 |
| TCGA-DU-6395 | 31 | male | oligoastrocytoma | G2 | Mutant | IDH1 | non-codel | 48.9867 | NaN | 1 |
| TCGA-DU-6397 | 45 | male | oligodendroglioma | G3 | Mutant | IDH1 | codel | 46.0297 | NaN | 3 |
| TCGA-DU-6399 | 54 | male | oligodendroglioma | G2 | Mutant | IDH1 | non-codel | 65.7098 | NaN | 1 |
| TCGA-DU-6400 | 66 | female | oligodendroglioma | G2 | Mutant | IDH1 | codel | 1.2156 | NaN | 3 |
| TCGA-DU-6401 | 31 | female | oligodendroglioma | G2 | Mutant | IDH1 | non-codel | 87.394 | NaN | 1 |
| TCGA-DU-6404 | 24 | female | oligodendroglioma | G3 | WT | N/A | non-codel | 133.6537 | 100 | 3 |
| TCGA-DU-6405 | 51 | female | astrocytoma | G3 | WT | N/A | non-codel | 19.8772 | 90 | 1 |
| TCGA-DU-6407 | 35 | female | oligodendroglioma | G2 | Mutant | IDH1 | non-codel | 94.4578 | 90 | 2 |
| TCGA-DU-6408 | 23 | female | oligodendroglioma | G3 | Mutant | IDH1 | non-codel | 114.0065 | 90 | 2 |
| TCGA-DU-6542 | 25 | male | oligoastrocytoma | G3 | Mutant | IDH1 | non-codel | 7.7209 | NaN | 1 |
| TCGA-DU-7008 | 41 | female | oligodendroglioma | G2 | Mutant | IDH1 | non-codel | 156.1265 | NaN | 3 |
| TCGA-DU-7010 | 58 | female | astrocytoma | G3 | Mutant | IDH1 | non-codel | 14.9818 | NaN | 2 |
| TCGA-DU-7015 | 41 | female | oligodendroglioma | G2 | Mutant | IDH1 | non-codel | 90.7124 | 90 | 1 |
| TCGA-DU-7018 | 57 | female | oligodendroglioma | G3 | Mutant | IDH1 | codel | 30.6536 | 90 | 2 |
| TCGA-DU-7019 | 39 | male | oligoastrocytoma | G3 | Mutant | IDH1 | non-codel | 26.2839 | 100 | 3 |
| TCGA-DU-7294 | 53 | female | oligodendroglioma | G2 | Mutant | IDH1 | codel | 94.2607 | 100 | 3 |
| TCGA-DU-7298 | 38 | female | astrocytoma | G3 | Mutant | IDH1 | non-codel | 18.9244 | 80 | 2 |
| TCGA-DU-7299 | 33 | male | astrocytoma | G3 | Mutant | IDH1 | non-codel | 43.9927 | 90 | 3 |
| TCGA-DU-7300 | 53 | female | oligodendroglioma | G3 | Mutant | IDH1 | codel | 61.9643 | 90 | 3 |
| TCGA-DU-7301 | 53 | male | oligodendroglioma | G2 | Mutant | IDH1 | non-codel | 25.8897 | 100 | 1 |
| TCGA-DU-7302 | 48 | female | oligodendroglioma | G3 | Mutant | IDH1 | codel | 60.2559 | 90 | 2 |
| TCGA-DU-7304 | 43 | male | oligoastrocytoma | G3 | Mutant | IDH1 | non-codel | 23.2941 | 80 | 2 |
| TCGA-DU-7306 | 67 | male | oligoastrocytoma | G2 | Mutant | IDH1 | non-codel | 41.9557 | 100 | 2 |
| TCGA-DU-7309 | 41 | female | oligodendroglioma | G3 | Mutant | IDH2 | non-codel | 2.7598 | 90 | 3 |
| TCGA-DU-8162 | 61 | female | oligoastrocytoma | G3 | WT | N/A | non-codel | 14.5876 | 80 | 2 |
| TCGA-DU-8163 | 29 | male | oligoastrocytoma | G3 | Mutant | IDH1 | non-codel | 20.6657 | 90 | 3 |
| TCGA-DU-8164 | 51 | male | oligodendroglioma | G2 | Mutant | IDH1 | codel | 21.3885 | NaN | 3 |
| TCGA-DU-8165 | 60 | female | oligodendroglioma | G3 | WT | N/A | non-codel | 19.1216 | 90 | 3 |
| TCGA-DU-8166 | 29 | female | oligoastrocytoma | G2 | Mutant | IDH1 | non-codel | 16.9531 | NaN | 3 |
| TCGA-DU-8167 | 69 | female | oligoastrocytoma | G2 | Mutant | IDH1 | non-codel | 15.4747 | 100 | 2 |
| TCGA-DU-8168 | 55 | female | oligodendroglioma | G3 | Mutant | IDH1 | codel | 14.1605 | 70 | 2 |
| TCGA-DU-A5TP | 33 | male | astrocytoma | G3 | Mutant | IDH1 | non-codel | 14.2262 | 70 | 1 |
| TCGA-DU-A5TR | 51 | male | oligoastrocytoma | G2 | Mutant | IDH1 | non-codel | 12.5834 | 90 | 1 |
| TCGA-DU-A5TS | 42 | male | oligodendroglioma | G2 | Mutant | IDH1 | non-codel | 15.1133 | 100 | 2 |
| TCGA-DU-A5TT | 70 | male | oligodendroglioma | G3 | WT | N/A | non-codel | 4.9611 | NaN | 1 |
| TCGA-DU-A5TU | 62 | female | astrocytoma | G2 | Mutant | IDH1 | non-codel | 3.6469 | 50 | 1 |
| TCGA-DU-A5TW | 33 | female | astrocytoma | G3 | Mutant | IDH1 | non-codel | 5.651 | 100 | 1 |
| TCGA-DU-A5TY | 46 | female | astrocytoma | G3 | WT | N/A | non-codel | 12.1892 | NaN | 2 |
| TCGA-DU-A6S2 | 37 | female | oligodendroglioma | G2 | Mutant | IDH1 | codel | 8.6408 | 70 | 3 |
| TCGA-DU-A6S3 | 60 | male | oligodendroglioma | G2 | Mutant | IDH1 | codel | 2.6941 | NaN | 3 |
| TCGA-DU-A6S6 | 35 | female | oligoastrocytoma | G2 | Mutant | IDH1 | codel | 77.9318 | 90 | 1 |
| TCGA-DU-A6S7 | 27 | female | astrocytoma | G3 | Mutant | IDH1 | non-codel | 7.2281 | 90 | 1 |
| TCGA-DU-A6S8 | 74 | female | oligodendroglioma | G3 | Mutant | IDH1 | codel | 5.9796 | 90 | 3 |
| TCGA-FG-5964 | 62 | male | oligodendroglioma | G2 | Mutant | IDH1 | codel | 34.3334 | NaN | 3 |
| TCGA-FG-6688 | 59 | female | astrocytoma | G3 | WT | N/A | non-codel | 18.7601 | 80 | 2 |
| TCGA-FG-6689 | 30 | male | astrocytoma | G2 | Mutant | IDH1 | non-codel | 14.9161 | 70 | 3 |
| TCGA-FG-6690 | 70 | male | oligodendroglioma | G2 | Mutant | IDH1 | non-codel | 24.9697 | 90 | 2 |
| TCGA-FG-6691 | 23 | female | astrocytoma | G2 | Mutant | IDH1 | non-codel | 23.9512 | 100 | 2 |
| TCGA-FG-6692 | 63 | male | oligodendroglioma | G3 | WT | N/A | non-codel | 18.4316 | NaN | 3 |
| TCGA-FG-7634 | 28 | male | oligodendroglioma | G2 | Mutant | IDH1 | codel | 15.3432 | NaN | 3 |
| TCGA-FG-7643 | 49 | female | oligoastrocytoma | G2 | WT | N/A | non-codel | 20.0743 | NaN | 2 |
| TCGA-FG-8189 | 33 | female | oligodendroglioma | G2 | Mutant | IDH2 | non-codel | 11.8606 | 70 | 2 |
| TCGA-FG-A4MT | 27 | female | oligodendroglioma | G2 | Mutant | IDH1 | non-codel | 38.2431 | 100 | 1 |
| TCGA-FG-A6IZ | 60 | male | oligodendroglioma | G2 | Mutant | IDH1 | codel | 4.2054 | NaN | 2 |
| TCGA-FG-A713 | 74 | female | oligoastrocytoma | G2 | Mutant | IDH1 | codel | 5.2239 | 60 | 3 |
| TCGA-HT-7473 | 28 | male | oligoastrocytoma | G2 | Mutant | IDH1 | non-codel | 16.526 | NaN | 1 |
| TCGA-HT-7475 | 67 | male | oligoastrocytoma | G3 | Mutant | IDH1 | non-codel | 17.4131 | 70 | 2 |
| TCGA-HT-7602 | 21 | male | oligodendroglioma | G2 | Mutant | IDH1 | non-codel | 29.8322 | NaN | 3 |
| TCGA-HT-7604 | 50 | male | astrocytoma | G2 | Mutant | IDH1 | non-codel | 107.8626 | NaN | 1 |
| TCGA-HT-7605 | 38 | male | oligodendroglioma | G2 | Mutant | IDH1 | codel | 4.5668 | NaN | 1 |
| TCGA-HT-7608 | 61 | male | oligoastrocytoma | G2 | Mutant | IDH1 | codel | 22.0456 | NaN | 3 |
| TCGA-HT-7616 | 75 | male | oligodendroglioma | G3 | Mutant | IDH1 | codel | 0.22998 | NaN | 3 |
| TCGA-HT-7680 | 32 | female | astrocytoma | G2 | WT | N/A | non-codel | 0.75566 | NaN | 3 |
| TCGA-HT-7686 | 29 | female | astrocytoma | G3 | Mutant | IDH1 | non-codel | 42.7114 | NaN | 3 |
| TCGA-HT-7690 | 29 | male | oligoastrocytoma | G3 | Mutant | IDH1 | non-codel | 0.098565 | NaN | 2 |
| TCGA-HT-7692 | 43 | male | oligoastrocytoma | G2 | Mutant | IDH1 | codel | 2.9569 | 100 | 2 |
| TCGA-HT-7693 | 51 | female | oligodendroglioma | G2 | Mutant | IDH1 | non-codel | 17.5117 | 90 | 1 |
| TCGA-HT-7694 | 60 | male | oligodendroglioma | G3 | Mutant | IDH1 | codel | 6.8995 | 90 | 3 |
| TCGA-HT-7855 | 39 | male | astrocytoma | G3 | Mutant | IDH1 | non-codel | 19.2201 | NaN | 2 |
| TCGA-HT-7856 | 35 | male | oligodendroglioma | G3 | Mutant | IDH2 | codel | 39.0645 | NaN | 2 |
| TCGA-HT-7860 | 60 | female | astrocytoma | G3 | WT | N/A | non-codel | 0.49282 | NaN | 2 |
| TCGA-HT-7874 | 41 | female | oligodendroglioma | G3 | Mutant | IDH1 | codel | 37.126 | NaN | 3 |
| TCGA-HT-7879 | 31 | male | oligoastrocytoma | G3 | Mutant | IDH1 | non-codel | 3.6797 | NaN | 3 |
| TCGA-HT-7882 | 66 | male | oligodendroglioma | G3 | WT | N/A | non-codel | 3.7126 | NaN | 1 |
| TCGA-HT-7884 | 44 | female | astrocytoma | G2 | Mutant | IDH1 | non-codel | 11.2692 | 80 | 1 |
| TCGA-HT-8018 | 40 | female | oligoastrocytoma | G2 | Mutant | IDH1 | non-codel | 21.4871 | NaN | 3 |
| TCGA-HT-8105 | 54 | male | oligodendroglioma | G3 | Mutant | IDH1 | codel | 6.2424 | NaN | 2 |
| TCGA-HT-8106 | 53 | male | astrocytoma | G3 | Mutant | IDH1 | non-codel | 0.098565 | NaN | 1 |
| TCGA-HT-8107 | 62 | male | oligodendroglioma | G2 | WT | N/A | non-codel | 0.45997 | NaN | 3 |
| TCGA-HT-8111 | 32 | male | oligoastrocytoma | G3 | Mutant | IDH1 | non-codel | 0.22998 | NaN | 3 |
| TCGA-HT-8113 | 49 | female | oligodendroglioma | G2 | Mutant | IDH2 | non-codel | 29.5694 | NaN | 1 |
| TCGA-HT-8114 | 36 | male | oligoastrocytoma | G3 | Mutant | IDH1 | non-codel | 3.8769 | NaN | 1 |
| TCGA-HT-8563 | 30 | female | astrocytoma | G3 | Mutant | IDH1 | non-codel | 16.0332 | NaN | 2 |
| TCGA-HT-A5RC | 70 | female | astrocytoma | G3 | WT | N/A | non-codel | 5.3225 | 40 | 3 |
| TCGA-HT-A61A | 20 | female | oligodendroglioma | G2 | Mutant | IDH1 | non-codel | 6.3739 | 80 | 1 |

Table 2: Subject wise 1p/19q co-deletion status and tumor histology

| **SUBJECT ID** | **Age** | **Gender** | **Histology** | **Grade** | **Data Collection** | **IDH**  **Status** | **IDH**  **Allele** | **1p/19q co-deletion** | **Cross-validation group** |
| --- | --- | --- | --- | --- | --- | --- | --- | --- | --- |
| TCGA-02-0003 | 50 | male | glioblastoma | G4 | HGG | WT | N/A | non-codel | 2 |
| TCGA-02-0006 | 56 | female | glioblastoma | G4 | HGG | WT | N/A | non-codel | 3 |
| TCGA-02-0009 | 61 | female | glioblastoma | G4 | HGG | WT | N/A | non-codel | 1 |
| TCGA-02-0011 | 18 | female | glioblastoma | G4 | HGG | WT | N/A | non-codel | 1 |
| TCGA-02-0027 | 33 | female | glioblastoma | G4 | HGG | WT | N/A | non-codel | 1 |
| TCGA-02-0033 | 54 | male | glioblastoma | G4 | HGG | WT | N/A | non-codel | 1 |
| TCGA-02-0034 | 60 | male | glioblastoma | G4 | HGG | WT | N/A | non-codel | 3 |
| TCGA-02-0037 | 74 | female | glioblastoma | G4 | HGG | WT | N/A | non-codel | 2 |
| TCGA-02-0046 | 61 | male | glioblastoma | G4 | HGG | WT | N/A | non-codel | 2 |
| TCGA-02-0047 | 78 | male | glioblastoma | G4 | HGG | WT | N/A | non-codel | 3 |
| TCGA-02-0048 | 80 | male | glioblastoma | G4 | HGG | WT | N/A | non-codel | 3 |
| TCGA-02-0054 | 44 | female | glioblastoma | G4 | HGG | WT | N/A | non-codel | 1 |
| TCGA-02-0060 | 66 | female | glioblastoma | G4 | HGG | WT | N/A | non-codel | 3 |
| TCGA-02-0064 | 50 | male | glioblastoma | G4 | HGG | WT | N/A | non-codel | 3 |
| TCGA-02-0068 | 57 | male | glioblastoma | G4 | HGG | WT | N/A | non-codel | 2 |
| TCGA-02-0069 | 31 | female | glioblastoma | G4 | HGG | WT | N/A | non-codel | 1 |
| TCGA-02-0070 | 70 | male | glioblastoma | G4 | HGG | WT | N/A | non-codel | 2 |
| TCGA-02-0075 | 63 | male | glioblastoma | G4 | HGG | WT | N/A | non-codel | 2 |
| TCGA-02-0085 | 63 | female | glioblastoma | G4 | HGG | WT | N/A | non-codel | 3 |
| TCGA-02-0086 | 45 | female | glioblastoma | G4 | HGG | WT | N/A | non-codel | 1 |
| TCGA-02-0102 | 42 | male | glioblastoma | G4 | HGG | WT | N/A | non-codel | 2 |
| TCGA-06-0119 | 81 | female | glioblastoma | G4 | HGG | WT | N/A | non-codel | 2 |
| TCGA-06-0122 | 84 | female | glioblastoma | G4 | HGG | WT | N/A | non-codel | 3 |
| TCGA-06-0127 | 67 | male | glioblastoma | G4 | HGG | WT | N/A | non-codel | 1 |
| TCGA-06-0128 | 66 | male | glioblastoma | G4 | HGG | Mutant | IDH1 | non-codel | 1 |
| TCGA-06-0129 | 30 | male | glioblastoma | G4 | HGG | Mutant | IDH1 | non-codel | 3 |
| TCGA-06-0132 | 49 | male | glioblastoma | G4 | HGG | WT | N/A | non-codel | 2 |
| TCGA-06-0133 | 64 | male | glioblastoma | G4 | HGG | WT | N/A | non-codel | 3 |
| TCGA-06-0137 | 63 | female | glioblastoma | G4 | HGG | WT | N/A | non-codel | 2 |
| TCGA-06-0138 | 43 | male | glioblastoma | G4 | HGG | WT | N/A | non-codel | 2 |
| TCGA-06-0139 | 40 | male | glioblastoma | G4 | HGG | WT | N/A | non-codel | 2 |
| TCGA-06-0142 | 81 | male | glioblastoma | G4 | HGG | WT | N/A | non-codel | 3 |
| TCGA-06-0143 | 58 | male | glioblastoma | G4 | HGG | WT | N/A | non-codel | 3 |
| TCGA-06-0145 | 53 | female | glioblastoma | G4 | HGG | WT | N/A | non-codel | 3 |
| TCGA-06-0147 | 51 | female | glioblastoma | G4 | HGG | WT | N/A | non-codel | 1 |
| TCGA-06-0154 | 54 | male | glioblastoma | G4 | HGG | WT | N/A | non-codel | 2 |
| TCGA-06-0157 | 63 | female | glioblastoma | G4 | HGG | WT | N/A | non-codel | 1 |
| TCGA-06-0158 | 73 | male | glioblastoma | G4 | HGG | WT | N/A | non-codel | 3 |
| TCGA-06-0166 | 51 | male | glioblastoma | G4 | HGG | WT | N/A | non-codel | 1 |
| TCGA-06-0168 | 59 | female | glioblastoma | G4 | HGG | WT | N/A | non-codel | 1 |
| TCGA-06-0174 | 54 | male | glioblastoma | G4 | HGG | WT | N/A | non-codel | 2 |
| TCGA-06-0176 | 34 | male | glioblastoma | G4 | HGG | WT | N/A | non-codel | 2 |
| TCGA-06-0184 | 63 | male | glioblastoma | G4 | HGG | WT | N/A | non-codel | 3 |
| TCGA-06-0185 | 54 | male | glioblastoma | G4 | HGG | WT | N/A | non-codel | 1 |
| TCGA-06-0187 | 69 | male | glioblastoma | G4 | HGG | WT | N/A | non-codel | 3 |
| TCGA-06-0188 | 71 | male | glioblastoma | G4 | HGG | WT | N/A | non-codel | 3 |
| TCGA-06-0189 | 55 | male | glioblastoma | G4 | HGG | WT | N/A | non-codel | 1 |
| TCGA-06-0190 | 62 | male | glioblastoma | G4 | HGG | WT | N/A | non-codel | 2 |
| TCGA-06-0192 | 58 | male | glioblastoma | G4 | HGG | WT | N/A | non-codel | 1 |
| TCGA-06-0213 | 55 | female | glioblastoma | G4 | HGG | WT | N/A | non-codel | 2 |
| TCGA-06-0237 | 75 | female | glioblastoma | G4 | HGG | WT | N/A | non-codel | 2 |
| TCGA-06-0238 | 46 | male | glioblastoma | G4 | HGG | WT | N/A | non-codel | 2 |
| TCGA-06-0241 | 65 | female | glioblastoma | G4 | HGG | WT | N/A | non-codel | 1 |
| TCGA-06-0644 | 71 | male | glioblastoma | G4 | HGG | WT | N/A | non-codel | 3 |
| TCGA-06-0645 | 55 | female | glioblastoma | G4 | HGG | WT | N/A | non-codel | 1 |
| TCGA-06-0646 | 60 | male | glioblastoma | G4 | HGG | WT | N/A | non-codel | 1 |
| TCGA-06-0648 | 77 | male | glioblastoma | G4 | HGG | WT | N/A | non-codel | 1 |
| TCGA-06-0649 | 73 | female | glioblastoma | G4 | HGG | WT | N/A | non-codel | 2 |
| TCGA-06-1806 | 47 | male | glioblastoma | G4 | HGG | WT | N/A | non-codel | 2 |
| TCGA-06-2570 | 21 | female | glioblastoma | G4 | HGG | Mutant | IDH1 | non-codel | 1 |
| TCGA-06-5408 | 54 | female | glioblastoma | G4 | HGG | WT | N/A | non-codel | 2 |
| TCGA-06-5412 | 78 | female | glioblastoma | G4 | HGG | WT | N/A | non-codel | 3 |
| TCGA-06-5413 | 67 | male | glioblastoma | G4 | HGG | WT | N/A | non-codel | 1 |
| TCGA-06-6389 | 49 | female | glioblastoma | G4 | HGG | Mutant | IDH1 | non-codel | 2 |
| TCGA-08-0390 | 69 | male | glioblastoma | G4 | HGG | WT | N/A | non-codel | 3 |
| TCGA-12-0616 | 36 | female | glioblastoma | G4 | HGG | WT | N/A | non-codel | 1 |
| TCGA-12-0829 | 75 | male | glioblastoma | G4 | HGG | WT | N/A | non-codel | 3 |
| TCGA-12-1093 | 66 | female | glioblastoma | G4 | HGG | WT | N/A | non-codel | 2 |
| TCGA-12-1598 | 75 | female | glioblastoma | G4 | HGG | WT | N/A | non-codel | 2 |
| TCGA-12-1602 | 58 | male | glioblastoma | G4 | HGG | WT | N/A | non-codel | 2 |
| TCGA-12-3650 | 46 | male | glioblastoma | G4 | HGG | WT | N/A | non-codel | 3 |
| TCGA-14-0789 | 54 | male | glioblastoma | G4 | HGG | WT | N/A | non-codel | 3 |
| TCGA-14-1456 | 23 | male | glioblastoma | G4 | HGG | Mutant | IDH1 | non-codel | 2 |
| TCGA-14-1794 | 59 | male | glioblastoma | G4 | HGG | WT | N/A | non-codel | 3 |
| TCGA-14-1829 | 57 | male | glioblastoma | G4 | HGG | WT | N/A | non-codel | 1 |
| TCGA-14-3477 | 38 | female | glioblastoma | G4 | HGG | WT | N/A | non-codel | 2 |
| TCGA-19-1388 | 58 | male | glioblastoma | G4 | HGG | WT | N/A | non-codel | 1 |
| TCGA-19-1390 | 63 | female | glioblastoma | G4 | HGG | WT | N/A | non-codel | 1 |
| TCGA-19-1789 | 69 | female | glioblastoma | G4 | HGG | WT | N/A | non-codel | 2 |
| TCGA-19-2624 | 51 | male | glioblastoma | G4 | HGG | WT | N/A | non-codel | 3 |
| TCGA-19-2631 | 74 | female | glioblastoma | G4 | HGG | WT | N/A | non-codel | 1 |
| TCGA-19-5954 | 72 | female | glioblastoma | G4 | HGG | WT | N/A | non-codel | 3 |
| TCGA-19-5958 | 56 | male | glioblastoma | G4 | HGG | WT | N/A | non-codel | 1 |
| TCGA-27-1835 | 53 | female | glioblastoma | G4 | HGG | WT | N/A | non-codel | 3 |
| TCGA-27-1838 | 59 | female | glioblastoma | G4 | HGG | WT | N/A | non-codel | 2 |
| TCGA-76-4926 | 68 | male | glioblastoma | G4 | HGG | WT | N/A | non-codel | 3 |
| TCGA-76-4934 | 66 | female | glioblastoma | G4 | HGG | WT | N/A | non-codel | 2 |
| TCGA-76-4935 | 52 | female | glioblastoma | G4 | HGG | WT | N/A | non-codel | 3 |
| TCGA-76-6191 | 57 | male | glioblastoma | G4 | HGG | WT | N/A | non-codel | 3 |
| TCGA-76-6192 | 74 | male | glioblastoma | G4 | HGG | WT | N/A | non-codel | 1 |
| TCGA-76-6193 | 78 | male | glioblastoma | G4 | HGG | WT | N/A | non-codel | 3 |
| TCGA-76-6280 | 57 | male | glioblastoma | G4 | HGG | WT | N/A | non-codel | 2 |
| TCGA-76-6282 | 63 | male | glioblastoma | G4 | HGG | WT | N/A | non-codel | 2 |
| TCGA-76-6285 | 64 | female | glioblastoma | G4 | HGG | WT | N/A | non-codel | 2 |
| TCGA-76-6656 | 66 | male | glioblastoma | G4 | HGG | WT | N/A | non-codel | 3 |
| TCGA-76-6657 | 74 | male | glioblastoma | G4 | HGG | WT | N/A | non-codel | 1 |
| TCGA-76-6661 | 54 | male | glioblastoma | G4 | HGG | WT | N/A | non-codel | 1 |
| TCGA-76-6662 | 58 | male | glioblastoma | G4 | HGG | WT | N/A | non-codel | 1 |
| TCGA-76-6663 | 44 | female | glioblastoma | G4 | HGG | WT | N/A | non-codel | 1 |
| TCGA-76-6664 | 49 | female | glioblastoma | G4 | HGG | WT | N/A | non-codel | 2 |
| TCGA-CS-4941 | 67 | male | astrocytoma | G3 | LGG | WT | N/A | non-codel | 2 |
| TCGA-CS-4942 | 44 | female | astrocytoma | G3 | LGG | Mutant | IDH1 | non-codel | 2 |
| TCGA-CS-4943 | 37 | male | astrocytoma | G3 | LGG | Mutant | IDH1 | non-codel | 2 |
| TCGA-CS-4944 | 50 | male | astrocytoma | G2 | LGG | Mutant | IDH1 | non-codel | 3 |
| TCGA-CS-5393 | 39 | male | astrocytoma | G3 | LGG | Mutant | IDH1 | non-codel | 3 |
| TCGA-CS-5395 | 43 | male | oligodendroglioma | G2 | LGG | WT | N/A | non-codel | 2 |
| TCGA-CS-5396 | 53 | female | oligodendroglioma | G3 | LGG | Mutant | IDH1 | codel | 2 |
| TCGA-CS-5397 | 54 | female | astrocytoma | G3 | LGG | WT | N/A | non-codel | 3 |
| TCGA-CS-6186 | 58 | male | oligoastrocytoma | G3 | LGG | WT | N/A | non-codel | 1 |
| TCGA-CS-6188 | 48 | male | astrocytoma | G3 | LGG | WT | N/A | non-codel | 1 |
| TCGA-CS-6290 | 31 | male | astrocytoma | G3 | LGG | Mutant | IDH1 | non-codel | 3 |
| TCGA-CS-6665 | 51 | female | astrocytoma | G3 | LGG | Mutant | IDH1 | non-codel | 2 |
| TCGA-CS-6666 | 22 | male | astrocytoma | G3 | LGG | Mutant | IDH1 | non-codel | 3 |
| TCGA-CS-6667 | 39 | female | astrocytoma | G2 | LGG | Mutant | IDH1 | non-codel | 1 |
| TCGA-CS-6668 | 57 | female | oligodendroglioma | G2 | LGG | Mutant | IDH1 | codel | 2 |
| TCGA-CS-6669 | 26 | female | oligodendroglioma | G2 | LGG | WT | N/A | non-codel | 1 |
| TCGA-DU-5849 | 48 | male | oligodendroglioma | G2 | LGG | Mutant | IDH1 | codel | 1 |
| TCGA-DU-5851 | 40 | female | oligoastrocytoma | G3 | LGG | Mutant | IDH1 | non-codel | 3 |
| TCGA-DU-5852 | 61 | female | oligoastrocytoma | G3 | LGG | WT | N/A | non-codel | 1 |
| TCGA-DU-5853 | 29 | male | oligoastrocytoma | G2 | LGG | Mutant | IDH1 | non-codel | 3 |
| TCGA-DU-5854 | 57 | female | astrocytoma | G3 | LGG | WT | N/A | non-codel | 2 |
| TCGA-DU-5855 | 49 | female | oligoastrocytoma | G3 | LGG | Mutant | IDH1 | non-codel | 1 |
| TCGA-DU-5871 | 37 | female | oligoastrocytoma | G2 | LGG | Mutant | IDH1 | non-codel | 2 |
| TCGA-DU-5872 | 43 | female | oligoastrocytoma | G2 | LGG | Mutant | IDH1 | non-codel | 1 |
| TCGA-DU-5874 | 62 | female | oligodendroglioma | G2 | LGG | Mutant | IDH1 | codel | 2 |
| TCGA-DU-6395 | 31 | male | oligoastrocytoma | G2 | LGG | Mutant | IDH1 | non-codel | 1 |
| TCGA-DU-6397 | 45 | male | oligodendroglioma | G3 | LGG | Mutant | IDH1 | codel | 2 |
| TCGA-DU-6399 | 54 | male | oligodendroglioma | G2 | LGG | Mutant | IDH1 | non-codel | 2 |
| TCGA-DU-6400 | 66 | female | oligodendroglioma | G2 | LGG | Mutant | IDH1 | codel | 1 |
| TCGA-DU-6401 | 31 | female | oligodendroglioma | G2 | LGG | Mutant | IDH1 | non-codel | 3 |
| TCGA-DU-6404 | 24 | female | oligodendroglioma | G3 | LGG | WT | N/A | non-codel | 3 |
| TCGA-DU-6405 | 51 | female | astrocytoma | G3 | LGG | WT | N/A | non-codel | 3 |
| TCGA-DU-6407 | 35 | female | oligodendroglioma | G2 | LGG | Mutant | IDH1 | non-codel | 3 |
| TCGA-DU-6408 | 23 | female | oligodendroglioma | G3 | LGG | Mutant | IDH1 | non-codel | 3 |
| TCGA-DU-7008 | 41 | female | oligodendroglioma | G2 | LGG | Mutant | IDH1 | non-codel | 2 |
| TCGA-DU-7010 | 58 | female | astrocytoma | G3 | LGG | Mutant | IDH1 | non-codel | 1 |
| TCGA-DU-7015 | 41 | female | oligodendroglioma | G2 | LGG | Mutant | IDH1 | non-codel | 3 |
| TCGA-DU-7018 | 57 | female | oligodendroglioma | G3 | LGG | Mutant | IDH1 | codel | 1 |
| TCGA-DU-7019 | 39 | male | oligoastrocytoma | G3 | LGG | Mutant | IDH1 | non-codel | 2 |
| TCGA-DU-7294 | 53 | female | oligodendroglioma | G2 | LGG | Mutant | IDH1 | codel | 2 |
| TCGA-DU-7298 | 38 | female | astrocytoma | G3 | LGG | Mutant | IDH1 | non-codel | 1 |
| TCGA-DU-7299 | 33 | male | astrocytoma | G3 | LGG | Mutant | IDH1 | non-codel | 1 |
| TCGA-DU-7300 | 53 | female | oligodendroglioma | G3 | LGG | Mutant | IDH1 | codel | 3 |
| TCGA-DU-7301 | 53 | male | oligodendroglioma | G2 | LGG | Mutant | IDH1 | non-codel | 2 |
| TCGA-DU-7302 | 48 | female | oligodendroglioma | G3 | LGG | Mutant | IDH1 | codel | 3 |
| TCGA-DU-7304 | 43 | male | oligoastrocytoma | G3 | LGG | Mutant | IDH1 | non-codel | 1 |
| TCGA-DU-7306 | 67 | male | oligoastrocytoma | G2 | LGG | Mutant | IDH1 | non-codel | 1 |
| TCGA-DU-7309 | 41 | female | oligodendroglioma | G3 | LGG | Mutant | IDH2 | non-codel | 3 |
| TCGA-DU-8162 | 61 | female | oligoastrocytoma | G3 | LGG | WT | N/A | non-codel | 2 |
| TCGA-DU-8163 | 29 | male | oligoastrocytoma | G3 | LGG | Mutant | IDH1 | non-codel | 3 |
| TCGA-DU-8164 | 51 | male | oligodendroglioma | G2 | LGG | Mutant | IDH1 | codel | 1 |
| TCGA-DU-8165 | 60 | female | oligodendroglioma | G3 | LGG | WT | N/A | non-codel | 3 |
| TCGA-DU-8166 | 29 | female | oligoastrocytoma | G2 | LGG | Mutant | IDH1 | non-codel | 2 |
| TCGA-DU-8167 | 69 | female | oligoastrocytoma | G2 | LGG | Mutant | IDH1 | non-codel | 2 |
| TCGA-DU-8168 | 55 | female | oligodendroglioma | G3 | LGG | Mutant | IDH1 | codel | 3 |
| TCGA-DU-A5TP | 33 | male | astrocytoma | G3 | LGG | Mutant | IDH1 | non-codel | 2 |
| TCGA-DU-A5TR | 51 | male | oligoastrocytoma | G2 | LGG | Mutant | IDH1 | non-codel | 2 |
| TCGA-DU-A5TS | 42 | male | oligodendroglioma | G2 | LGG | Mutant | IDH1 | non-codel | 1 |
| TCGA-DU-A5TT | 70 | male | oligodendroglioma | G3 | LGG | WT | N/A | non-codel | 2 |
| TCGA-DU-A5TU | 62 | female | astrocytoma | G2 | LGG | Mutant | IDH1 | non-codel | 2 |
| TCGA-DU-A5TW | 33 | female | astrocytoma | G3 | LGG | Mutant | IDH1 | non-codel | 2 |
| TCGA-DU-A5TY | 46 | female | astrocytoma | G3 | LGG | WT | N/A | non-codel | 2 |
| TCGA-DU-A6S2 | 37 | female | oligodendroglioma | G2 | LGG | Mutant | IDH1 | codel | 2 |
| TCGA-DU-A6S3 | 60 | male | oligodendroglioma | G2 | LGG | Mutant | IDH1 | codel | 3 |
| TCGA-DU-A6S6 | 35 | female | oligoastrocytoma | G2 | LGG | Mutant | IDH1 | codel | 1 |
| TCGA-DU-A6S7 | 27 | female | astrocytoma | G3 | LGG | Mutant | IDH1 | non-codel | 3 |
| TCGA-DU-A6S8 | 74 | female | oligodendroglioma | G3 | LGG | Mutant | IDH1 | codel | 1 |
| TCGA-FG-5964 | 62 | male | oligodendroglioma | G2 | LGG | Mutant | IDH1 | codel | 2 |
| TCGA-FG-6688 | 59 | female | astrocytoma | G3 | LGG | WT | N/A | non-codel | 1 |
| TCGA-FG-6689 | 30 | male | astrocytoma | G2 | LGG | Mutant | IDH1 | non-codel | 1 |
| TCGA-FG-6690 | 70 | male | oligodendroglioma | G2 | LGG | Mutant | IDH1 | non-codel | 3 |
| TCGA-FG-6691 | 23 | female | astrocytoma | G2 | LGG | Mutant | IDH1 | non-codel | 1 |
| TCGA-FG-6692 | 63 | male | oligodendroglioma | G3 | LGG | WT | N/A | non-codel | 3 |
| TCGA-FG-7634 | 28 | male | oligodendroglioma | G2 | LGG | Mutant | IDH1 | codel | 2 |
| TCGA-FG-7643 | 49 | female | oligoastrocytoma | G2 | LGG | WT | N/A | non-codel | 1 |
| TCGA-FG-8189 | 33 | female | oligodendroglioma | G2 | LGG | Mutant | IDH2 | non-codel | 2 |
| TCGA-FG-A4MT | 27 | female | oligodendroglioma | G2 | LGG | Mutant | IDH1 | non-codel | 2 |
| TCGA-FG-A6IZ | 60 | male | oligodendroglioma | G2 | LGG | Mutant | IDH1 | codel | 3 |
| TCGA-FG-A713 | 74 | female | oligoastrocytoma | G2 | LGG | Mutant | IDH1 | codel | 1 |
| TCGA-HT-7473 | 28 | male | oligoastrocytoma | G2 | LGG | Mutant | IDH1 | non-codel | 3 |
| TCGA-HT-7475 | 67 | male | oligoastrocytoma | G3 | LGG | Mutant | IDH1 | non-codel | 3 |
| TCGA-HT-7602 | 21 | male | oligodendroglioma | G2 | LGG | Mutant | IDH1 | non-codel | 3 |
| TCGA-HT-7604 | 50 | male | astrocytoma | G2 | LGG | Mutant | IDH1 | non-codel | 1 |
| TCGA-HT-7605 | 38 | male | oligodendroglioma | G2 | LGG | Mutant | IDH1 | codel | 3 |
| TCGA-HT-7608 | 61 | male | oligoastrocytoma | G2 | LGG | Mutant | IDH1 | codel | 3 |
| TCGA-HT-7616 | 75 | male | oligodendroglioma | G3 | LGG | Mutant | IDH1 | codel | 2 |
| TCGA-HT-7680 | 32 | female | astrocytoma | G2 | LGG | WT | N/A | non-codel | 1 |
| TCGA-HT-7686 | 29 | female | astrocytoma | G3 | LGG | Mutant | IDH1 | non-codel | 2 |
| TCGA-HT-7690 | 29 | male | oligoastrocytoma | G3 | LGG | Mutant | IDH1 | non-codel | 3 |
| TCGA-HT-7692 | 43 | male | oligoastrocytoma | G2 | LGG | Mutant | IDH1 | codel | 3 |
| TCGA-HT-7693 | 51 | female | oligodendroglioma | G2 | LGG | Mutant | IDH1 | non-codel | 1 |
| TCGA-HT-7694 | 60 | male | oligodendroglioma | G3 | LGG | Mutant | IDH1 | codel | 1 |
| TCGA-HT-7855 | 39 | male | astrocytoma | G3 | LGG | Mutant | IDH1 | non-codel | 1 |
| TCGA-HT-7856 | 35 | male | oligodendroglioma | G3 | LGG | Mutant | IDH2 | codel | 1 |
| TCGA-HT-7860 | 60 | female | astrocytoma | G3 | LGG | WT | N/A | non-codel | 1 |
| TCGA-HT-7874 | 41 | female | oligodendroglioma | G3 | LGG | Mutant | IDH1 | codel | 1 |
| TCGA-HT-7879 | 31 | male | oligoastrocytoma | G3 | LGG | Mutant | IDH1 | non-codel | 3 |
| TCGA-HT-7882 | 66 | male | oligodendroglioma | G3 | LGG | WT | N/A | non-codel | 2 |
| TCGA-HT-7884 | 44 | female | astrocytoma | G2 | LGG | Mutant | IDH1 | non-codel | 1 |
| TCGA-HT-8018 | 40 | female | oligoastrocytoma | G2 | LGG | Mutant | IDH1 | non-codel | 1 |
| TCGA-HT-8105 | 54 | male | oligodendroglioma | G3 | LGG | Mutant | IDH1 | codel | 3 |
| TCGA-HT-8106 | 53 | male | astrocytoma | G3 | LGG | Mutant | IDH1 | non-codel | 3 |
| TCGA-HT-8107 | 62 | male | oligodendroglioma | G2 | LGG | WT | N/A | non-codel | 3 |
| TCGA-HT-8111 | 32 | male | oligoastrocytoma | G3 | LGG | Mutant | IDH1 | non-codel | 2 |
| TCGA-HT-8113 | 49 | female | oligodendroglioma | G2 | LGG | Mutant | IDH2 | non-codel | 2 |
| TCGA-HT-8114 | 36 | male | oligoastrocytoma | G3 | LGG | Mutant | IDH1 | non-codel | 2 |
| TCGA-HT-8563 | 30 | female | astrocytoma | G3 | LGG | Mutant | IDH1 | non-codel | 3 |
| TCGA-HT-A5RC | 70 | female | astrocytoma | G3 | LGG | WT | N/A | non-codel | 3 |
| TCGA-HT-A61A | 20 | female | oligodendroglioma | G2 | LGG | Mutant | IDH1 | non-codel | 1 |
| LGG-104 | NaN | NaN | Oligoastrocytoma | G3 | LGG | NaN | NaN | codel | 1 |
| LGG-203 | NaN | NaN | Astrocytoma | G3 | LGG | NaN | NaN | non-codel | 1 |
| LGG-210 | NaN | NaN | Oligoastrocytoma | G2 | LGG | NaN | NaN | non-codel | 1 |
| LGG-216 | NaN | NaN | Oligoastrocytoma | G2 | LGG | NaN | NaN | codel | 2 |
| LGG-218 | NaN | NaN | Oligodendroglioma | G2 | LGG | NaN | NaN | codel | 1 |
| LGG-219 | NaN | NaN | Astrocytoma | G3 | LGG | NaN | NaN | non-codel | 3 |
| LGG-220 | NaN | NaN | Oligodendroglioma | G2 | LGG | NaN | NaN | codel | 1 |
| LGG-223 | NaN | NaN | Oligodendroglioma | G3 | LGG | NaN | NaN | codel | 1 |
| LGG-225 | NaN | NaN | Oligoastrocytoma | G2 | LGG | NaN | NaN | codel | 2 |
| LGG-229 | NaN | NaN | Oligodendroglioma | G2 | LGG | NaN | NaN | codel | 2 |
| LGG-231 | NaN | NaN | Oligoastrocytoma | G3 | LGG | NaN | NaN | codel | 1 |
| LGG-233 | NaN | NaN | Oligodendroglioma | G2 | LGG | NaN | NaN | codel | 3 |
| LGG-234 | NaN | NaN | Oligodendroglioma | G3 | LGG | NaN | NaN | non-codel | 3 |
| LGG-240 | NaN | NaN | Astrocytoma | G3 | LGG | NaN | NaN | non-codel | 3 |
| LGG-241 | NaN | NaN | Oligoastrocytoma | G2 | LGG | NaN | NaN | non-codel | 2 |
| LGG-246 | NaN | NaN | Oligoastrocytoma | G3 | LGG | NaN | NaN | codel | 3 |
| LGG-249 | NaN | NaN | Oligodendroglioma | G2 | LGG | NaN | NaN | codel | 3 |
| LGG-254 | NaN | NaN | Oligodendroglioma | G3 | LGG | NaN | NaN | codel | 2 |
| LGG-260 | NaN | NaN | Oligodendroglioma | G3 | LGG | NaN | NaN | codel | 1 |
| LGG-261 | NaN | NaN | Oligodendroglioma | G2 | LGG | NaN | NaN | codel | 1 |
| LGG-263 | NaN | NaN | Oligoastrocytoma | G3 | LGG | NaN | NaN | non-codel | 3 |
| LGG-269 | NaN | NaN | Oligodendroglioma | G3 | LGG | NaN | NaN | codel | 3 |
| LGG-273 | NaN | NaN | Oligoastrocytoma | G3 | LGG | NaN | NaN | non-codel | 1 |
| LGG-274 | NaN | NaN | Oligoastrocytoma | G2 | LGG | NaN | NaN | codel | 2 |
| LGG-277 | NaN | NaN | Astrocytoma | G2 | LGG | NaN | NaN | non-codel | 3 |
| LGG-278 | NaN | NaN | Oligoastrocytoma | G2 | LGG | NaN | NaN | codel | 2 |
| LGG-280 | NaN | NaN | Oligoastrocytoma | G2 | LGG | NaN | NaN | non-codel | 3 |
| LGG-282 | NaN | NaN | Oligoastrocytoma | G3 | LGG | NaN | NaN | codel | 2 |
| LGG-285 | NaN | NaN | Oligoastrocytoma | G2 | LGG | NaN | NaN | non-codel | 2 |
| LGG-286 | NaN | NaN | Oligodendroglioma | G2 | LGG | NaN | NaN | non-codel | 1 |
| LGG-288 | NaN | NaN | Oligoastrocytoma | G3 | LGG | NaN | NaN | codel | 1 |
| LGG-289 | NaN | NaN | Oligodendroglioma | G2 | LGG | NaN | NaN | codel | 2 |
| LGG-293 | NaN | NaN | Oligoastrocytoma | G2 | LGG | NaN | NaN | non-codel | 3 |
| LGG-295 | NaN | NaN | Oligoastrocytoma | G3 | LGG | NaN | NaN | codel | 3 |
| LGG-296 | NaN | NaN | Oligodendroglioma | G2 | LGG | NaN | NaN | codel | 2 |
| LGG-297 | NaN | NaN | Oligoastrocytoma | G2 | LGG | NaN | NaN | non-codel | 3 |
| LGG-298 | NaN | NaN | Oligoastrocytoma | G3 | LGG | NaN | NaN | codel | 3 |
| LGG-303 | NaN | NaN | Oligodendroglioma | G2 | LGG | NaN | NaN | codel | 1 |
| LGG-304 | NaN | NaN | Oligodendroglioma | G2 | LGG | NaN | NaN | codel | 3 |
| LGG-305 | NaN | NaN | Oligodendroglioma | G2 | LGG | NaN | NaN | codel | 3 |
| LGG-306 | NaN | NaN | Astrocytoma | G3 | LGG | NaN | NaN | non-codel | 3 |
| LGG-307 | NaN | NaN | Oligoastrocytoma | G3 | LGG | NaN | NaN | codel | 2 |
| LGG-308 | NaN | NaN | Oligodendroglioma | G2 | LGG | NaN | NaN | codel | 3 |
| LGG-310 | NaN | NaN | Oligoastrocytoma | G2 | LGG | NaN | NaN | codel | 2 |
| LGG-311 | NaN | NaN | Astrocytoma | G2 | LGG | NaN | NaN | non-codel | 3 |
| LGG-313 | NaN | NaN | Oligoastrocytoma | G2 | LGG | NaN | NaN | non-codel | 2 |
| LGG-314 | NaN | NaN | Oligoastrocytoma | G2 | LGG | NaN | NaN | non-codel | 1 |
| LGG-315 | NaN | NaN | Oligodendroglioma | G2 | LGG | NaN | NaN | codel | 2 |
| LGG-316 | NaN | NaN | Oligodendroglioma | G3 | LGG | NaN | NaN | codel | 1 |
| LGG-320 | NaN | NaN | Oligoastrocytoma | G2 | LGG | NaN | NaN | codel | 1 |
| LGG-321 | NaN | NaN | Oligoastrocytoma | G2 | LGG | NaN | NaN | non-codel | 2 |
| LGG-325 | NaN | NaN | Oligoastrocytoma | G2 | LGG | NaN | NaN | codel | 3 |
| LGG-326 | NaN | NaN | Oligoastrocytoma | G2 | LGG | NaN | NaN | codel | 1 |
| LGG-327 | NaN | NaN | Oligoastrocytoma | G2 | LGG | NaN | NaN | non-codel | 3 |
| LGG-330 | NaN | NaN | Oligoastrocytoma | G3 | LGG | NaN | NaN | codel | 3 |
| LGG-331 | NaN | NaN | Oligodendroglioma | G2 | LGG | NaN | NaN | codel | 2 |
| LGG-333 | NaN | NaN | Oligoastrocytoma | G3 | LGG | NaN | NaN | codel | 2 |
| LGG-334 | NaN | NaN | Oligoastrocytoma | G2 | LGG | NaN | NaN | non-codel | 2 |
| LGG-337 | NaN | NaN | Oligoastrocytoma | G2 | LGG | NaN | NaN | codel | 1 |
| LGG-338 | NaN | NaN | Oligoastrocytoma | G3 | LGG | NaN | NaN | non-codel | 3 |
| LGG-341 | NaN | NaN | Oligoastrocytoma | G3 | LGG | NaN | NaN | codel | 2 |
| LGG-343 | NaN | NaN | Oligoastrocytoma | G2 | LGG | NaN | NaN | non-codel | 2 |
| LGG-344 | NaN | NaN | Oligodendroglioma | G2 | LGG | NaN | NaN | codel | 1 |
| LGG-345 | NaN | NaN | Oligoastrocytoma | G3 | LGG | NaN | NaN | codel | 3 |
| LGG-346 | NaN | NaN | Astrocytoma | G3 | LGG | NaN | NaN | non-codel | 2 |
| LGG-348 | NaN | NaN | Oligoastrocytoma | G2 | LGG | NaN | NaN | codel | 1 |
| LGG-350 | NaN | NaN | Oligoastrocytoma | G2 | LGG | NaN | NaN | codel | 1 |
| LGG-351 | NaN | NaN | Oligoastrocytoma | G2 | LGG | NaN | NaN | non-codel | 1 |
| LGG-352 | NaN | NaN | Oligodendroglioma | G2 | LGG | NaN | NaN | codel | 2 |
| LGG-354 | NaN | NaN | Oligoastrocytoma | G3 | LGG | NaN | NaN | non-codel | 1 |
| LGG-355 | NaN | NaN | Astrocytoma | G3 | LGG | NaN | NaN | codel | 1 |
| LGG-357 | NaN | NaN | Oligoastrocytoma | G2 | LGG | NaN | NaN | codel | 1 |
| LGG-359 | NaN | NaN | Oligodendroglioma | G2 | LGG | NaN | NaN | codel | 3 |
| LGG-360 | NaN | NaN | Oligodendroglioma | G2 | LGG | NaN | NaN | codel | 3 |
| LGG-361 | NaN | NaN | Oligoastrocytoma | G2 | LGG | NaN | NaN | codel | 1 |
| LGG-363 | NaN | NaN | Oligoastrocytoma | G2 | LGG | NaN | NaN | non-codel | 2 |
| LGG-365 | NaN | NaN | Oligoastrocytoma | G3 | LGG | NaN | NaN | codel | 3 |
| LGG-367 | NaN | NaN | Oligoastrocytoma | G2 | LGG | NaN | NaN | codel | 3 |
| LGG-371 | NaN | NaN | Astrocytoma | G3 | LGG | NaN | NaN | non-codel | 1 |
| LGG-373 | NaN | NaN | Oligoastrocytoma | G3 | LGG | NaN | NaN | codel | 2 |
| LGG-374 | NaN | NaN | Oligoastrocytoma | G2 | LGG | NaN | NaN | non-codel | 3 |
| LGG-375 | NaN | NaN | Oligoastrocytoma | G2 | LGG | NaN | NaN | non-codel | 2 |
| LGG-377 | NaN | NaN | Oligodendroglioma | G3 | LGG | NaN | NaN | codel | 1 |
| LGG-380 | NaN | NaN | Oligoastrocytoma | G3 | LGG | NaN | NaN | codel | 1 |
| LGG-383 | NaN | NaN | Oligoastrocytoma | G2 | LGG | NaN | NaN | codel | 3 |
| LGG-385 | NaN | NaN | Oligoastrocytoma | G3 | LGG | NaN | NaN | codel | 2 |
| LGG-387 | NaN | NaN | Oligoastrocytoma | G3 | LGG | NaN | NaN | codel | 2 |
| LGG-388 | NaN | NaN | Oligoastrocytoma | G2 | LGG | NaN | NaN | codel | 1 |
| LGG-391 | NaN | NaN | Oligoastrocytoma | G2 | LGG | NaN | NaN | non-codel | 1 |
| LGG-394 | NaN | NaN | Oligodendroglioma | G3 | LGG | NaN | NaN | codel | 3 |
| LGG-395 | NaN | NaN | Oligoastrocytoma | G3 | LGG | NaN | NaN | codel | 1 |
| LGG-396 | NaN | NaN | Oligoastrocytoma | G2 | LGG | NaN | NaN | codel | 1 |
| LGG-492 | NaN | NaN | Oligoastrocytoma | G2 | LGG | NaN | NaN | codel | 2 |
| LGG-500 | NaN | NaN | Oligoastrocytoma | G2 | LGG | NaN | NaN | non-codel | 1 |
| LGG-506 | NaN | NaN | Oligoastrocytoma | G2 | LGG | NaN | NaN | non-codel | 1 |
| LGG-515 | NaN | NaN | Oligoastrocytoma | G2 | LGG | NaN | NaN | codel | 3 |
| LGG-516 | NaN | NaN | Oligoastrocytoma | G3 | LGG | NaN | NaN | non-codel | 2 |
| LGG-518 | NaN | NaN | Astrocytoma | G3 | LGG | NaN | NaN | non-codel | 3 |
| LGG-519 | NaN | NaN | Oligoastrocytoma | G2 | LGG | NaN | NaN | non-codel | 1 |
| LGG-520 | NaN | NaN | Oligodendroglioma | G2 | LGG | NaN | NaN | codel | 2 |
| LGG-525 | NaN | NaN | Oligodendroglioma | G2 | LGG | NaN | NaN | codel | 2 |
| LGG-527 | NaN | NaN | Oligoastrocytoma | G2 | LGG | NaN | NaN | codel | 3 |
| LGG-532 | NaN | NaN | Oligoastrocytoma | G3 | LGG | NaN | NaN | non-codel | 3 |
| LGG-533 | NaN | NaN | Oligoastrocytoma | G2 | LGG | NaN | NaN | non-codel | 1 |
| LGG-537 | NaN | NaN | Oligoastrocytoma | G2 | LGG | NaN | NaN | non-codel | 2 |
| LGG-545 | NaN | NaN | Oligoastrocytoma | G2 | LGG | NaN | NaN | non-codel | 1 |
| LGG-547 | NaN | NaN | Oligodendroglioma | G2 | LGG | NaN | NaN | codel | 2 |
| LGG-550 | NaN | NaN | Oligoastrocytoma | G3 | LGG | NaN | NaN | codel | 3 |
| LGG-552 | NaN | NaN | Astrocytoma | G3 | LGG | NaN | NaN | non-codel | 2 |
| LGG-558 | NaN | NaN | Oligoastrocytoma | G2 | LGG | NaN | NaN | non-codel | 1 |
| LGG-561 | NaN | NaN | Oligoastrocytoma | G3 | LGG | NaN | NaN | codel | 2 |
| LGG-563 | NaN | NaN | Oligodendroglioma | G2 | LGG | NaN | NaN | codel | 2 |
| LGG-565 | NaN | NaN | Oligoastrocytoma | G2 | LGG | NaN | NaN | codel | 3 |
| LGG-566 | NaN | NaN | Oligoastrocytoma | G2 | LGG | NaN | NaN | codel | 3 |
| LGG-570 | NaN | NaN | Oligoastrocytoma | G2 | LGG | NaN | NaN | codel | 2 |
| LGG-572 | NaN | NaN | Astrocytoma | G3 | LGG | NaN | NaN | codel | 3 |
| LGG-573 | NaN | NaN | Oligoastrocytoma | G2 | LGG | NaN | NaN | codel | 1 |
| LGG-574 | NaN | NaN | Oligodendroglioma | G2 | LGG | NaN | NaN | non-codel | 1 |
| LGG-576 | NaN | NaN | Oligoastrocytoma | G2 | LGG | NaN | NaN | codel | 1 |
| LGG-579 | NaN | NaN | Oligoastrocytoma | G2 | LGG | NaN | NaN | codel | 1 |
| LGG-581 | NaN | NaN | Oligoastrocytoma | G2 | LGG | NaN | NaN | codel | 2 |
| LGG-582 | NaN | NaN | Oligodendroglioma | G2 | LGG | NaN | NaN | codel | 2 |
| LGG-585 | NaN | NaN | Astrocytoma | G2 | LGG | NaN | NaN | non-codel | 3 |
| LGG-587 | NaN | NaN | Oligodendroglioma | G3 | LGG | NaN | NaN | codel | 3 |
| LGG-589 | NaN | NaN | Oligoastrocytoma | G2 | LGG | NaN | NaN | non-codel | 2 |
| LGG-590 | NaN | NaN | Oligoastrocytoma | G2 | LGG | NaN | NaN | codel | 1 |
| LGG-591 | NaN | NaN | Astrocytoma | G3 | LGG | NaN | NaN | non-codel | 1 |
| LGG-593 | NaN | NaN | Oligodendroglioma | G3 | LGG | NaN | NaN | codel | 1 |
| LGG-594 | NaN | NaN | Oligoastrocytoma | G3 | LGG | NaN | NaN | non-codel | 2 |
| LGG-597 | NaN | NaN | Oligoastrocytoma | G2 | LGG | NaN | NaN | codel | 1 |
| LGG-600 | NaN | NaN | Oligoastrocytoma | G3 | LGG | NaN | NaN | codel | 3 |
| LGG-601 | NaN | NaN | Astrocytoma | G3 | LGG | NaN | NaN | non-codel | 3 |
| LGG-604 | NaN | NaN | Oligoastrocytoma | G2 | LGG | NaN | NaN | codel | 2 |
| LGG-607 | NaN | NaN | Oligodendroglioma | G2 | LGG | NaN | NaN | codel | 3 |
| LGG-609 | NaN | NaN | Oligoastrocytoma | G2 | LGG | NaN | NaN | non-codel | 2 |
| LGG-610 | NaN | NaN | Oligoastrocytoma | G2 | LGG | NaN | NaN | non-codel | 3 |
| LGG-612 | NaN | NaN | Oligodendroglioma | G2 | LGG | NaN | NaN | codel | 2 |
| LGG-613 | NaN | NaN | Oligoastrocytoma | G2 | LGG | NaN | NaN | non-codel | 1 |
| LGG-614 | NaN | NaN | Oligodendroglioma | G2 | LGG | NaN | NaN | codel | 3 |
| LGG-616 | NaN | NaN | Oligoastrocytoma | G2 | LGG | NaN | NaN | codel | 3 |
| LGG-620 | NaN | NaN | Oligodendroglioma | G2 | LGG | NaN | NaN | codel | 3 |
| LGG-622 | NaN | NaN | Oligoastrocytoma | G2 | LGG | NaN | NaN | non-codel | 2 |
| LGG-624 | NaN | NaN | Oligoastrocytoma | G3 | LGG | NaN | NaN | non-codel | 2 |
| LGG-625 | NaN | NaN | Oligoastrocytoma | G2 | LGG | NaN | NaN | non-codel | 2 |
| LGG-626 | NaN | NaN | Astrocytoma | G3 | LGG | NaN | NaN | codel | 2 |
| LGG-630 | NaN | NaN | Oligodendroglioma | G2 | LGG | NaN | NaN | codel | 3 |
| LGG-631 | NaN | NaN | Oligoastrocytoma | G2 | LGG | NaN | NaN | non-codel | 1 |
| LGG-632 | NaN | NaN | Oligoastrocytoma | G2 | LGG | NaN | NaN | codel | 2 |
| LGG-634 | NaN | NaN | Oligodendroglioma | G3 | LGG | NaN | NaN | codel | 2 |
| LGG-637 | NaN | NaN | Oligodendroglioma | G2 | LGG | NaN | NaN | codel | 1 |
| LGG-639 | NaN | NaN | Oligoastrocytoma | G2 | LGG | NaN | NaN | codel | 1 |
| LGG-642 | NaN | NaN | Oligodendroglioma | G3 | LGG | NaN | NaN | codel | 2 |
| LGG-647 | NaN | NaN | Oligoastrocytoma | G2 | LGG | NaN | NaN | non-codel | 3 |
| LGG-648 | NaN | NaN | Astrocytoma | G2 | LGG | NaN | NaN | codel | 3 |
| LGG-651 | NaN | NaN | Oligodendroglioma | G2 | LGG | NaN | NaN | codel | 3 |
| LGG-658 | NaN | NaN | Oligodendroglioma | G3 | LGG | NaN | NaN | codel | 3 |
| LGG-659 | NaN | NaN | Oligoastrocytoma | G2 | LGG | NaN | NaN | codel | 2 |
| LGG-660 | NaN | NaN | Oligoastrocytoma | G2 | LGG | NaN | NaN | codel | 1 |
| LGG-766 | NaN | NaN | Oligoastrocytoma | G2 | LGG | NaN | NaN | non-codel | 1 |

Table 3: Subject wise MGMT status and tumor histology

| **Subject ID** | **Age** | **Gender** | **Histology** | **Grade** | **TCGA Data Collection** | **IDH mutation Status** | **1p/19q co-deletion status** | **MGMT promoter status** | **Cross-validation group** |
| --- | --- | --- | --- | --- | --- | --- | --- | --- | --- |
| TCGA-02-0003 | 50 | male | glioblastoma | G4 | TCGA-GBM | WT | non-codel | Unmethylated | 2 |
| TCGA-02-0006 | 56 | female | glioblastoma | G4 | TCGA-GBM | WT | non-codel | Unmethylated | 2 |
| TCGA-02-0009 | 61 | female | glioblastoma | G4 | TCGA-GBM | WT | non-codel | Unmethylated | 2 |
| TCGA-02-0011 | 18 | female | glioblastoma | G4 | TCGA-GBM | WT | non-codel | Methylated | 2 |
| TCGA-02-0027 | 33 | female | glioblastoma | G4 | TCGA-GBM | WT | non-codel | Unmethylated | 2 |
| TCGA-02-0033 | 54 | male | glioblastoma | G4 | TCGA-GBM | WT | non-codel | Methylated | 1 |
| TCGA-02-0034 | 60 | male | glioblastoma | G4 | TCGA-GBM | WT | non-codel | Unmethylated | 1 |
| TCGA-02-0037 | 74 | female | glioblastoma | G4 | TCGA-GBM | WT | non-codel | Unmethylated | 1 |
| TCGA-02-0046 | 61 | male | glioblastoma | G4 | TCGA-GBM | WT | non-codel | Methylated | 1 |
| TCGA-02-0047 | 78 | male | glioblastoma | G4 | TCGA-GBM | WT | non-codel | Unmethylated | 1 |
| TCGA-02-0060 | 66 | female | glioblastoma | G4 | TCGA-GBM | WT | non-codel | Methylated | 2 |
| TCGA-02-0064 | 50 | male | glioblastoma | G4 | TCGA-GBM | WT | non-codel | Methylated | 3 |
| TCGA-02-0069 | 31 | female | glioblastoma | G4 | TCGA-GBM | WT | non-codel | Methylated | 2 |
| TCGA-02-0075 | 63 | male | glioblastoma | G4 | TCGA-GBM | WT | non-codel | Methylated | 2 |
| TCGA-02-0086 | 45 | female | glioblastoma | G4 | TCGA-GBM | WT | non-codel | Unmethylated | 2 |
| TCGA-02-0102 | 42 | male | glioblastoma | G4 | TCGA-GBM | WT | non-codel | Unmethylated | 1 |
| TCGA-06-0119 | 81 | female | glioblastoma | G4 | TCGA-GBM | WT | non-codel | Methylated | 2 |
| TCGA-06-0122 | 84 | female | glioblastoma | G4 | TCGA-GBM | WT | non-codel | Unmethylated | 2 |
| TCGA-06-0128 | 66 | male | glioblastoma | G4 | TCGA-GBM | Mutant | non-codel | Methylated | 3 |
| TCGA-06-0129 | 30 | male | glioblastoma | G4 | TCGA-GBM | Mutant | non-codel | Methylated | 1 |
| TCGA-06-0133 | 64 | male | glioblastoma | G4 | TCGA-GBM | WT | non-codel | Unmethylated | 2 |
| TCGA-06-0137 | 63 | female | glioblastoma | G4 | TCGA-GBM | WT | non-codel | Unmethylated | 2 |
| TCGA-06-0142 | 81 | male | glioblastoma | G4 | TCGA-GBM | WT | non-codel | Unmethylated | 1 |
| TCGA-06-0143 | 58 | male | glioblastoma | G4 | TCGA-GBM | WT | non-codel | Unmethylated | 1 |
| TCGA-06-0145 | 53 | female | glioblastoma | G4 | TCGA-GBM | WT | non-codel | Methylated | 1 |
| TCGA-06-0147 | 51 | female | glioblastoma | G4 | TCGA-GBM | WT | non-codel | Methylated | 2 |
| TCGA-06-0148 | 76 | male | glioblastoma | G4 | TCGA-GBM | WT | non-codel | Unmethylated | 1 |
| TCGA-06-0881 | 50 | male | glioblastoma | G4 | TCGA-GBM | WT | non-codel | Unmethylated | 1 |
| TCGA-06-1806 | 47 | male | glioblastoma | G4 | TCGA-GBM | WT | non-codel | Unmethylated | 2 |
| TCGA-06-2570 | 21 | female | glioblastoma | G4 | TCGA-GBM | Mutant | non-codel | Methylated | 1 |
| TCGA-06-5408 | 54 | female | glioblastoma | G4 | TCGA-GBM | WT | non-codel | Unmethylated | 3 |
| TCGA-06-5412 | 78 | female | glioblastoma | G4 | TCGA-GBM | WT | non-codel | Methylated | 2 |
| TCGA-06-5413 | 67 | male | glioblastoma | G4 | TCGA-GBM | WT | non-codel | Unmethylated | 1 |
| TCGA-06-5417 | 45 | female | glioblastoma | G4 | TCGA-GBM | Mutant | NA | Methylated | 3 |
| TCGA-06-6389 | 49 | female | glioblastoma | G4 | TCGA-GBM | Mutant | non-codel | Methylated | 3 |
| TCGA-12-0829 | 75 | male | glioblastoma | G4 | TCGA-GBM | WT | non-codel | Methylated | 2 |
| TCGA-12-1093 | 66 | female | glioblastoma | G4 | TCGA-GBM | WT | non-codel | Unmethylated | 1 |
| TCGA-12-1598 | 75 | female | glioblastoma | G4 | TCGA-GBM | WT | non-codel | Methylated | 3 |
| TCGA-12-1601 | NaN | NA | NA | NA | TCGA-GBM | WT | NA | Unmethylated | 2 |
| TCGA-12-1602 | 58 | male | glioblastoma | G4 | TCGA-GBM | WT | non-codel | Methylated | 2 |
| TCGA-12-3650 | 46 | male | glioblastoma | G4 | TCGA-GBM | WT | non-codel | Unmethylated | 3 |
| TCGA-14-0789 | 54 | male | glioblastoma | G4 | TCGA-GBM | WT | non-codel | Methylated | 2 |
| TCGA-14-1456 | 23 | male | glioblastoma | G4 | TCGA-GBM | Mutant | non-codel | Unmethylated | 3 |
| TCGA-14-1794 | 59 | male | glioblastoma | G4 | TCGA-GBM | WT | non-codel | Unmethylated | 2 |
| TCGA-14-1829 | 57 | male | glioblastoma | G4 | TCGA-GBM | WT | non-codel | Unmethylated | 2 |
| TCGA-14-3477 | 38 | female | glioblastoma | G4 | TCGA-GBM | WT | non-codel | Unmethylated | 3 |
| TCGA-19-1390 | 63 | female | glioblastoma | G4 | TCGA-GBM | WT | non-codel | Methylated | 1 |
| TCGA-19-1789 | 69 | female | glioblastoma | G4 | TCGA-GBM | WT | non-codel | Methylated | 2 |
| TCGA-19-1791 | 82 | female | glioblastoma | G4 | TCGA-GBM | WT | non-codel | Unmethylated | 1 |
| TCGA-19-2620 | 70 | male | glioblastoma | G4 | TCGA-GBM | WT | non-codel | Methylated | 3 |
| TCGA-19-2624 | 51 | male | glioblastoma | G4 | TCGA-GBM | WT | non-codel | Unmethylated | 1 |
| TCGA-19-2631 | 74 | female | glioblastoma | G4 | TCGA-GBM | WT | non-codel | Methylated | 3 |
| TCGA-19-5953 | 58 | male | glioblastoma | G4 | TCGA-GBM | WT | non-codel | Methylated | 1 |
| TCGA-19-5954 | 72 | female | glioblastoma | G4 | TCGA-GBM | WT | non-codel | Methylated | 3 |
| TCGA-19-5958 | 56 | male | glioblastoma | G4 | TCGA-GBM | WT | non-codel | Unmethylated | 1 |
| TCGA-27-1830 | 57 | male | glioblastoma | G4 | TCGA-GBM | WT | non-codel | Unmethylated | 2 |
| TCGA-27-1835 | 53 | female | glioblastoma | G4 | TCGA-GBM | WT | non-codel | Methylated | 1 |
| TCGA-27-1836 | 33 | female | glioblastoma | G4 | TCGA-GBM | WT | non-codel | Methylated | 1 |
| TCGA-27-1838 | 59 | female | glioblastoma | G4 | TCGA-GBM | WT | non-codel | Unmethylated | 3 |
| TCGA-76-4925 | 76 | male | glioblastoma | G4 | TCGA-GBM | WT | non-codel | Methylated | 2 |
| TCGA-76-4926 | 68 | male | glioblastoma | G4 | TCGA-GBM | WT | non-codel | Unmethylated | 3 |
| TCGA-76-4927 | 58 | male | glioblastoma | G4 | TCGA-GBM | WT | NA | Unmethylated | 2 |
| TCGA-76-4928 | 85 | female | glioblastoma | G4 | TCGA-GBM | WT | non-codel | Methylated | 1 |
| TCGA-76-4929 | 76 | female | glioblastoma | G4 | TCGA-GBM | WT | non-codel | Methylated | 3 |
| TCGA-76-4931 | 70 | female | glioblastoma | G4 | TCGA-GBM | WT | non-codel | Unmethylated | 3 |
| TCGA-76-4932 | 50 | female | glioblastoma | G4 | TCGA-GBM | WT | NA | Methylated | 3 |
| TCGA-76-4934 | 66 | female | glioblastoma | G4 | TCGA-GBM | WT | non-codel | Methylated | 1 |
| TCGA-76-4935 | 52 | female | glioblastoma | G4 | TCGA-GBM | WT | non-codel | Methylated | 2 |
| TCGA-76-6191 | 57 | male | glioblastoma | G4 | TCGA-GBM | WT | non-codel | Unmethylated | 3 |
| TCGA-76-6192 | 74 | male | glioblastoma | G4 | TCGA-GBM | WT | non-codel | Unmethylated | 2 |
| TCGA-76-6193 | 78 | male | glioblastoma | G4 | TCGA-GBM | WT | non-codel | Unmethylated | 2 |
| TCGA-76-6280 | 57 | male | glioblastoma | G4 | TCGA-GBM | WT | non-codel | Methylated | 1 |
| TCGA-76-6282 | 63 | male | glioblastoma | G4 | TCGA-GBM | WT | non-codel | Unmethylated | 2 |
| TCGA-76-6285 | 64 | female | glioblastoma | G4 | TCGA-GBM | WT | non-codel | Unmethylated | 3 |
| TCGA-76-6286 | 60 | male | glioblastoma | G4 | TCGA-GBM | WT | non-codel | Unmethylated | 1 |
| TCGA-76-6656 | 66 | male | glioblastoma | G4 | TCGA-GBM | WT | non-codel | Methylated | 3 |
| TCGA-76-6657 | 74 | male | glioblastoma | G4 | TCGA-GBM | WT | non-codel | Methylated | 2 |
| TCGA-76-6661 | 54 | male | glioblastoma | G4 | TCGA-GBM | WT | non-codel | Unmethylated | 3 |
| TCGA-76-6662 | 58 | male | glioblastoma | G4 | TCGA-GBM | WT | non-codel | Unmethylated | 1 |
| TCGA-76-6663 | 44 | female | glioblastoma | G4 | TCGA-GBM | WT | non-codel | Unmethylated | 3 |
| TCGA-76-6664 | 49 | female | glioblastoma | G4 | TCGA-GBM | WT | non-codel | Methylated | 3 |
| TCGA-CS-4938 | 31 | female | astrocytoma | G2 | TCGA-LGG | Mutant | non-codel | Unmethylated | 3 |
| TCGA-CS-4941 | 67 | male | astrocytoma | G3 | TCGA-LGG | WT | non-codel | Methylated | 1 |
| TCGA-CS-4942 | 44 | female | astrocytoma | G3 | TCGA-LGG | Mutant | non-codel | Unmethylated | 1 |
| TCGA-CS-4943 | 37 | male | astrocytoma | G3 | TCGA-LGG | Mutant | non-codel | Methylated | 3 |
| TCGA-CS-4944 | 50 | male | astrocytoma | G2 | TCGA-LGG | Mutant | non-codel | Methylated | 2 |
| TCGA-CS-5390 | 47 | female | oligodendroglioma | G2 | TCGA-LGG | Mutant | codel | Methylated | 2 |
| TCGA-CS-5393 | 39 | male | astrocytoma | G3 | TCGA-LGG | Mutant | non-codel | Methylated | 2 |
| TCGA-CS-5394 | 40 | male | astrocytoma | G3 | TCGA-LGG | Mutant | non-codel | Methylated | 1 |
| TCGA-CS-5395 | 43 | male | oligodendroglioma | G2 | TCGA-LGG | WT | non-codel | Unmethylated | 1 |
| TCGA-CS-5396 | 53 | female | oligodendroglioma | G3 | TCGA-LGG | Mutant | codel | Methylated | 3 |
| TCGA-CS-5397 | 54 | female | astrocytoma | G3 | TCGA-LGG | WT | non-codel | Unmethylated | 2 |
| TCGA-CS-6186 | 58 | male | oligoastrocytoma | G3 | TCGA-LGG | WT | non-codel | Unmethylated | 1 |
| TCGA-CS-6188 | 48 | male | astrocytoma | G3 | TCGA-LGG | WT | non-codel | Unmethylated | 2 |
| TCGA-CS-6290 | 31 | male | astrocytoma | G3 | TCGA-LGG | Mutant | non-codel | Methylated | 1 |
| TCGA-CS-6665 | 51 | female | astrocytoma | G3 | TCGA-LGG | Mutant | non-codel | Methylated | 3 |
| TCGA-CS-6666 | 22 | male | astrocytoma | G3 | TCGA-LGG | Mutant | non-codel | Methylated | 2 |
| TCGA-CS-6667 | 39 | female | astrocytoma | G2 | TCGA-LGG | Mutant | non-codel | Methylated | 1 |
| TCGA-CS-6668 | 57 | female | oligodendroglioma | G2 | TCGA-LGG | Mutant | codel | Methylated | 1 |
| TCGA-CS-6669 | 26 | female | oligodendroglioma | G2 | TCGA-LGG | WT | non-codel | Unmethylated | 3 |
| TCGA-DU-5849 | 48 | male | oligodendroglioma | G2 | TCGA-LGG | Mutant | codel | Methylated | 1 |
| TCGA-DU-5851 | 40 | female | oligoastrocytoma | G3 | TCGA-LGG | Mutant | non-codel | Unmethylated | 3 |
| TCGA-DU-5852 | 61 | female | oligoastrocytoma | G3 | TCGA-LGG | WT | non-codel | Methylated | 3 |
| TCGA-DU-5853 | 29 | male | oligoastrocytoma | G2 | TCGA-LGG | Mutant | non-codel | Methylated | 2 |
| TCGA-DU-5854 | 57 | female | astrocytoma | G3 | TCGA-LGG | WT | non-codel | Unmethylated | 2 |
| TCGA-DU-5855 | 49 | female | oligoastrocytoma | G3 | TCGA-LGG | Mutant | non-codel | Methylated | 2 |
| TCGA-DU-5871 | 37 | female | oligoastrocytoma | G2 | TCGA-LGG | Mutant | non-codel | Methylated | 3 |
| TCGA-DU-5872 | 43 | female | oligoastrocytoma | G2 | TCGA-LGG | Mutant | non-codel | Methylated | 2 |
| TCGA-DU-5874 | 62 | female | oligodendroglioma | G2 | TCGA-LGG | Mutant | codel | Methylated | 3 |
| TCGA-DU-6395 | 31 | male | oligoastrocytoma | G2 | TCGA-LGG | Mutant | non-codel | Methylated | 1 |
| TCGA-DU-6397 | 45 | male | oligodendroglioma | G3 | TCGA-LGG | Mutant | codel | Methylated | 2 |
| TCGA-DU-6399 | 54 | male | oligodendroglioma | G2 | TCGA-LGG | Mutant | non-codel | Methylated | 1 |
| TCGA-DU-6400 | 66 | female | oligodendroglioma | G2 | TCGA-LGG | Mutant | codel | Methylated | 3 |
| TCGA-DU-6401 | 31 | female | oligodendroglioma | G2 | TCGA-LGG | Mutant | non-codel | Methylated | 3 |
| TCGA-DU-6404 | 24 | female | oligodendroglioma | G3 | TCGA-LGG | WT | non-codel | Unmethylated | 3 |
| TCGA-DU-6405 | 51 | female | astrocytoma | G3 | TCGA-LGG | WT | non-codel | Methylated | 3 |
| TCGA-DU-6407 | 35 | female | oligodendroglioma | G2 | TCGA-LGG | Mutant | non-codel | Methylated | 3 |
| TCGA-DU-6408 | 23 | female | oligodendroglioma | G3 | TCGA-LGG | Mutant | non-codel | Methylated | 1 |
| TCGA-DU-7008 | 41 | female | oligodendroglioma | G2 | TCGA-LGG | Mutant | non-codel | Methylated | 1 |
| TCGA-DU-7010 | 58 | female | astrocytoma | G3 | TCGA-LGG | Mutant | non-codel | Methylated | 3 |
| TCGA-DU-7013 | 59 | male | astrocytoma | G3 | TCGA-LGG | WT | non-codel | Unmethylated | 1 |
| TCGA-DU-7015 | 41 | female | oligodendroglioma | G2 | TCGA-LGG | Mutant | non-codel | Methylated | 2 |
| TCGA-DU-7018 | 57 | female | oligodendroglioma | G3 | TCGA-LGG | Mutant | codel | Methylated | 1 |
| TCGA-DU-7019 | 39 | male | oligoastrocytoma | G3 | TCGA-LGG | Mutant | non-codel | Methylated | 1 |
| TCGA-DU-7294 | 53 | female | oligodendroglioma | G2 | TCGA-LGG | Mutant | codel | Methylated | 1 |
| TCGA-DU-7298 | 38 | female | astrocytoma | G3 | TCGA-LGG | Mutant | non-codel | Methylated | 2 |
| TCGA-DU-7299 | 33 | male | astrocytoma | G3 | TCGA-LGG | Mutant | non-codel | Methylated | 3 |
| TCGA-DU-7300 | 53 | female | oligodendroglioma | G3 | TCGA-LGG | Mutant | codel | Methylated | 3 |
| TCGA-DU-7301 | 53 | male | oligodendroglioma | G2 | TCGA-LGG | Mutant | non-codel | Methylated | 3 |
| TCGA-DU-7302 | 48 | female | oligodendroglioma | G3 | TCGA-LGG | Mutant | codel | Methylated | 1 |
| TCGA-DU-7304 | 43 | male | oligoastrocytoma | G3 | TCGA-LGG | Mutant | non-codel | Methylated | 3 |
| TCGA-DU-7306 | 67 | male | oligoastrocytoma | G2 | TCGA-LGG | Mutant | non-codel | Methylated | 1 |
| TCGA-DU-7309 | 41 | female | oligodendroglioma | G3 | TCGA-LGG | Mutant | non-codel | Methylated | 3 |
| TCGA-DU-8158 | 57 | female | astrocytoma | G3 | TCGA-LGG | WT | non-codel | Unmethylated | 3 |
| TCGA-DU-8162 | 61 | female | oligoastrocytoma | G3 | TCGA-LGG | WT | non-codel | Unmethylated | 1 |
| TCGA-DU-8164 | 51 | male | oligodendroglioma | G2 | TCGA-LGG | Mutant | codel | Methylated | 1 |
| TCGA-DU-8165 | 60 | female | oligodendroglioma | G3 | TCGA-LGG | WT | non-codel | Unmethylated | 1 |
| TCGA-DU-8166 | 29 | female | oligoastrocytoma | G2 | TCGA-LGG | Mutant | non-codel | Methylated | 2 |
| TCGA-DU-8167 | 69 | female | oligoastrocytoma | G2 | TCGA-LGG | Mutant | non-codel | Methylated | 3 |
| TCGA-DU-8168 | 55 | female | oligodendroglioma | G3 | TCGA-LGG | Mutant | codel | Methylated | 3 |
| TCGA-DU-A5TP | 33 | male | astrocytoma | G3 | TCGA-LGG | Mutant | non-codel | Methylated | 2 |
| TCGA-DU-A5TR | 51 | male | oligoastrocytoma | G2 | TCGA-LGG | Mutant | non-codel | Methylated | 2 |
| TCGA-DU-A5TS | 42 | male | oligodendroglioma | G2 | TCGA-LGG | Mutant | non-codel | Methylated | 2 |
| TCGA-DU-A5TT | 70 | male | oligodendroglioma | G3 | TCGA-LGG | WT | non-codel | Methylated | 2 |
| TCGA-DU-A5TU | 62 | female | astrocytoma | G2 | TCGA-LGG | Mutant | non-codel | Methylated | 1 |
| TCGA-DU-A5TW | 33 | female | astrocytoma | G3 | TCGA-LGG | Mutant | non-codel | Methylated | 2 |
| TCGA-DU-A5TY | 46 | female | astrocytoma | G3 | TCGA-LGG | WT | non-codel | Methylated | 2 |
| TCGA-DU-A6S2 | 37 | female | oligodendroglioma | G2 | TCGA-LGG | Mutant | codel | Methylated | 2 |
| TCGA-DU-A6S3 | 60 | male | oligodendroglioma | G2 | TCGA-LGG | Mutant | codel | Methylated | 3 |
| TCGA-DU-A6S6 | 35 | female | oligoastrocytoma | G2 | TCGA-LGG | Mutant | codel | Methylated | 1 |
| TCGA-DU-A6S7 | 27 | female | astrocytoma | G3 | TCGA-LGG | Mutant | non-codel | Methylated | 2 |
| TCGA-DU-A6S8 | 74 | female | oligodendroglioma | G3 | TCGA-LGG | Mutant | codel | Methylated | 2 |
| TCGA-FG-5963 | 23 | male | astrocytoma | G3 | TCGA-LGG | WT | non-codel | Unmethylated | 2 |
| TCGA-FG-5964 | 62 | male | oligodendroglioma | G2 | TCGA-LGG | Mutant | codel | Methylated | 1 |
| TCGA-FG-6688 | 59 | female | astrocytoma | G3 | TCGA-LGG | WT | non-codel | Methylated | 2 |
| TCGA-FG-6689 | 30 | male | astrocytoma | G2 | TCGA-LGG | Mutant | non-codel | Methylated | 3 |
| TCGA-FG-6690 | 70 | male | oligodendroglioma | G2 | TCGA-LGG | Mutant | non-codel | Methylated | 2 |
| TCGA-FG-6691 | 23 | female | astrocytoma | G2 | TCGA-LGG | Mutant | non-codel | Unmethylated | 3 |
| TCGA-FG-6692 | 63 | male | oligodendroglioma | G3 | TCGA-LGG | WT | non-codel | Methylated | 1 |
| TCGA-FG-7634 | 28 | male | oligodendroglioma | G2 | TCGA-LGG | Mutant | codel | Methylated | 1 |
| TCGA-FG-7637 | 49 | male | oligoastrocytoma | G2 | TCGA-LGG | Mutant | non-codel | Methylated | 1 |
| TCGA-FG-8189 | 33 | female | oligodendroglioma | G2 | TCGA-LGG | Mutant | non-codel | Methylated | 3 |
| TCGA-FG-A4MT | 27 | female | oligodendroglioma | G2 | TCGA-LGG | Mutant | non-codel | Methylated | 2 |
| TCGA-FG-A4MU | 58 | male | oligoastrocytoma | G3 | TCGA-LGG | WT | non-codel | Methylated | 1 |
| TCGA-FG-A6IZ | 60 | male | oligodendroglioma | G2 | TCGA-LGG | Mutant | codel | Methylated | 3 |
| TCGA-FG-A6J1 | 44 | female | oligodendroglioma | G2 | TCGA-LGG | Mutant | codel | Methylated | 2 |
| TCGA-FG-A713 | 74 | female | oligoastrocytoma | G2 | TCGA-LGG | Mutant | codel | Methylated | 3 |
| TCGA-FG-A87N | 37 | female | astrocytoma | G3 | TCGA-LGG | Mutant | non-codel | Methylated | 3 |
| TCGA-HT-7468 | 30 | male | oligodendroglioma | G3 | TCGA-LGG | Mutant | codel | Methylated | 1 |
| TCGA-HT-7469 | 30 | male | oligodendroglioma | G3 | TCGA-LGG | WT | non-codel | Methylated | 1 |
| TCGA-HT-7471 | 37 | female | oligodendroglioma | G3 | TCGA-LGG | Mutant | codel | Methylated | 2 |
| TCGA-HT-7472 | 38 | male | oligodendroglioma | G2 | TCGA-LGG | Mutant | non-codel | Methylated | 1 |
| TCGA-HT-7473 | 28 | male | oligoastrocytoma | G2 | TCGA-LGG | Mutant | non-codel | Unmethylated | 3 |
| TCGA-HT-7475 | 67 | male | oligoastrocytoma | G3 | TCGA-LGG | Mutant | non-codel | Methylated | 3 |
| TCGA-HT-7476 | 26 | male | astrocytoma | G2 | TCGA-LGG | Mutant | non-codel | Methylated | 1 |
| TCGA-HT-7478 | 36 | male | astrocytoma | G2 | TCGA-LGG | Mutant | non-codel | Unmethylated | 1 |
| TCGA-HT-7481 | 39 | male | oligodendroglioma | G2 | TCGA-LGG | Mutant | codel | Methylated | 2 |
| TCGA-HT-7602 | 21 | male | oligodendroglioma | G2 | TCGA-LGG | Mutant | non-codel | Methylated | 2 |
| TCGA-HT-7603 | 29 | male | oligodendroglioma | G2 | TCGA-LGG | Mutant | non-codel | Methylated | 2 |
| TCGA-HT-7605 | 38 | male | oligodendroglioma | G2 | TCGA-LGG | Mutant | codel | Methylated | 3 |
| TCGA-HT-7606 | 30 | female | astrocytoma | G2 | TCGA-LGG | Mutant | non-codel | Unmethylated | 1 |
| TCGA-HT-7608 | 61 | male | oligoastrocytoma | G2 | TCGA-LGG | Mutant | codel | Methylated | 2 |
| TCGA-HT-7616 | 75 | male | oligodendroglioma | G3 | TCGA-LGG | Mutant | codel | Methylated | 3 |
| TCGA-HT-7680 | 32 | female | astrocytoma | G2 | TCGA-LGG | WT | non-codel | Unmethylated | 3 |
| TCGA-HT-7684 | 58 | male | oligoastrocytoma | G3 | TCGA-LGG | Mutant | non-codel | Methylated | 3 |
| TCGA-HT-7686 | 29 | female | astrocytoma | G3 | TCGA-LGG | Mutant | non-codel | Methylated | 3 |
| TCGA-HT-7690 | 29 | male | oligoastrocytoma | G3 | TCGA-LGG | Mutant | non-codel | Methylated | 3 |
| TCGA-HT-7692 | 43 | male | oligoastrocytoma | G2 | TCGA-LGG | Mutant | codel | Methylated | 1 |
| TCGA-HT-7693 | 51 | female | oligodendroglioma | G2 | TCGA-LGG | Mutant | non-codel | Methylated | 1 |
| TCGA-HT-7694 | 60 | male | oligodendroglioma | G3 | TCGA-LGG | Mutant | codel | Methylated | 2 |
| TCGA-HT-7695 | 29 | female | oligodendroglioma | G2 | TCGA-LGG | Mutant | codel | Methylated | 1 |
| TCGA-HT-7854 | 62 | male | astrocytoma | G2 | TCGA-LGG | WT | non-codel | Unmethylated | 3 |
| TCGA-HT-7855 | 39 | male | astrocytoma | G3 | TCGA-LGG | Mutant | non-codel | Methylated | 2 |
| TCGA-HT-7856 | 35 | male | oligodendroglioma | G3 | TCGA-LGG | Mutant | codel | Methylated | 2 |
| TCGA-HT-7860 | 60 | female | astrocytoma | G3 | TCGA-LGG | WT | non-codel | Methylated | 2 |
| TCGA-HT-7874 | 41 | female | oligodendroglioma | G3 | TCGA-LGG | Mutant | codel | Methylated | 1 |
| TCGA-HT-7877 | 20 | female | oligodendroglioma | G2 | TCGA-LGG | Mutant | codel | Methylated | 3 |
| TCGA-HT-7879 | 31 | male | oligoastrocytoma | G3 | TCGA-LGG | Mutant | non-codel | Methylated | 1 |
| TCGA-HT-7880 | 30 | male | oligoastrocytoma | G2 | TCGA-LGG | Mutant | non-codel | Methylated | 3 |
| TCGA-HT-7882 | 66 | male | oligodendroglioma | G3 | TCGA-LGG | WT | non-codel | Methylated | 2 |
| TCGA-HT-7884 | 44 | female | astrocytoma | G2 | TCGA-LGG | Mutant | non-codel | Methylated | 1 |
| TCGA-HT-7902 | 30 | female | oligoastrocytoma | G2 | TCGA-LGG | Mutant | non-codel | Methylated | 3 |
| TCGA-HT-8010 | 64 | female | oligodendroglioma | G2 | TCGA-LGG | Mutant | codel | Methylated | 3 |
| TCGA-HT-8013 | 37 | female | oligoastrocytoma | G2 | TCGA-LGG | Mutant | non-codel | Methylated | 1 |
| TCGA-HT-8015 | 21 | male | astrocytoma | G2 | TCGA-LGG | WT | non-codel | Unmethylated | 1 |
| TCGA-HT-8018 | 40 | female | oligoastrocytoma | G2 | TCGA-LGG | Mutant | non-codel | Methylated | 3 |
| TCGA-HT-8019 | 34 | female | oligodendroglioma | G3 | TCGA-LGG | WT | non-codel | Unmethylated | 1 |
| TCGA-HT-8105 | 54 | male | oligodendroglioma | G3 | TCGA-LGG | Mutant | codel | Methylated | 3 |
| TCGA-HT-8106 | 53 | male | astrocytoma | G3 | TCGA-LGG | Mutant | non-codel | Methylated | 1 |
| TCGA-HT-8107 | 62 | male | oligodendroglioma | G2 | TCGA-LGG | WT | non-codel | Methylated | 2 |
| TCGA-HT-8108 | 26 | female | oligodendroglioma | G2 | TCGA-LGG | Mutant | non-codel | Methylated | 1 |
| TCGA-HT-8111 | 32 | male | oligoastrocytoma | G3 | TCGA-LGG | Mutant | non-codel | Methylated | 2 |
| TCGA-HT-8113 | 49 | female | oligodendroglioma | G2 | TCGA-LGG | Mutant | non-codel | Methylated | 3 |
| TCGA-HT-8114 | 36 | male | oligoastrocytoma | G3 | TCGA-LGG | Mutant | non-codel | Methylated | 2 |
| TCGA-HT-8558 | 29 | female | oligodendroglioma | G2 | TCGA-LGG | WT | non-codel | Unmethylated | 2 |
| TCGA-HT-8563 | 30 | female | astrocytoma | G3 | TCGA-LGG | Mutant | non-codel | Unmethylated | 3 |
| TCGA-HT-8564 | 47 | male | astrocytoma | G3 | TCGA-LGG | WT | non-codel | Unmethylated | 2 |
| TCGA-HT-A4DS | 55 | female | astrocytoma | G3 | TCGA-LGG | WT | non-codel | Unmethylated | 2 |
| TCGA-HT-A5R5 | 33 | female | oligodendroglioma | G2 | TCGA-LGG | Mutant | non-codel | Methylated | 2 |
| TCGA-HT-A5RB | 24 | male | astrocytoma | G2 | TCGA-LGG | Mutant | non-codel | Methylated | 1 |
| TCGA-HT-A5RC | 70 | female | astrocytoma | G3 | TCGA-LGG | WT | non-codel | Unmethylated | 2 |
| TCGA-HT-A616 | 36 | female | astrocytoma | G2 | TCGA-LGG | Mutant | non-codel | Methylated | 3 |
| TCGA-HT-A61A | 20 | female | oligodendroglioma | G2 | TCGA-LGG | Mutant | non-codel | Methylated | 3 |
| TCGA-HT-A61B | NaN | NaN | NaN | NaN | TCGA-LGG | Mutant | non-codel | Methylated | 2 |
| W1_19961025 | NaN | NaN | NaN | NaN | NaN | NaN | NaN | Unmethylated | 2 |
| W10_19970429 | NaN | NaN | NaN | NaN | NaN | NaN | NaN | Methylated | 1 |
| W12_19970620 | NaN | NaN | NaN | NaN | NaN | NaN | NaN | Unmethylated | 3 |
| W13_19970822 | NaN | NaN | NaN | NaN | NaN | NaN | NaN | Unmethylated | 3 |
| W16_19971015 | NaN | NaN | NaN | NaN | NaN | NaN | NaN | Unmethylated | 1 |
| W18_19971110 | NaN | NaN | NaN | NaN | NaN | NaN | NaN | Methylated | 1 |
| W2_19961101 | NaN | NaN | NaN | NaN | NaN | NaN | NaN | Methylated | 3 |
| W20_19970516 | NaN | NaN | NaN | NaN | NaN | NaN | NaN | Unmethylated | 1 |
| W21_19980105 | NaN | NaN | NaN | NaN | NaN | NaN | NaN | Unmethylated | 2 |
| W22_19980102 | NaN | NaN | NaN | NaN | NaN | NaN | NaN | Methylated | 1 |
| W29_19980521 | NaN | NaN | NaN | NaN | NaN | NaN | NaN | Unmethylated | 3 |
| W30_19980608 | NaN | NaN | NaN | NaN | NaN | NaN | NaN | Methylated | 2 |
| W31_19980629 | NaN | NaN | NaN | NaN | NaN | NaN | NaN | Unmethylated | 3 |
| W32_19980701 | NaN | NaN | NaN | NaN | NaN | NaN | NaN | Methylated | 3 |
| W33_19980704 | NaN | NaN | NaN | NaN | NaN | NaN | NaN | Methylated | 3 |
| W34_19980713 | NaN | NaN | NaN | NaN | NaN | NaN | NaN | Unmethylated | 2 |
| W36_19980714 | NaN | NaN | NaN | NaN | NaN | NaN | NaN | Unmethylated | 1 |
| W38_19980910 | NaN | NaN | NaN | NaN | NaN | NaN | NaN | Methylated | 2 |
| W39_19980919 | NaN | NaN | NaN | NaN | NaN | NaN | NaN | Methylated | 1 |
| W5_19961211 | NaN | NaN | NaN | NaN | NaN | NaN | NaN | Unmethylated | 3 |
| W54_20000902 | NaN | NaN | NaN | NaN | NaN | NaN | NaN | Unmethylated | 3 |
| W7_19961218 | NaN | NaN | NaN | NaN | NaN | NaN | NaN | Methylated | 1 |
| W9_19970410 | NaN | NaN | NaN | NaN | NaN | NaN | NaN | Unmethylated | 3 |

Table 4. Cross-validation data distribution for Combined Dataset.

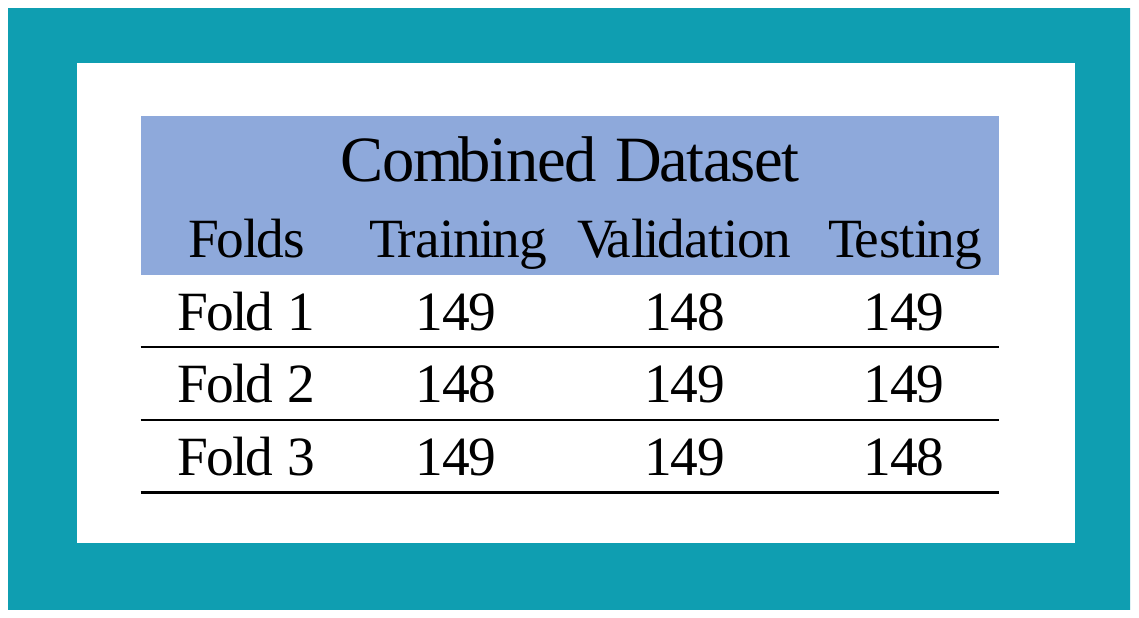
